# Decoding calcium oscillation frequency in transcriptional regulation

**DOI:** 10.1101/2025.10.10.676024

**Authors:** Parvaneh Nikpour, Manuel Varas-Godoy, Per Uhlén, Erik Smedler

## Abstract

Cells continuously experience fluctuating intracellular calcium (Ca²⁺) signals that orchestrate diverse processes such as transcription, proliferation, and apoptosis. Temporal features of Ca²⁺ dynamics, including oscillation frequency, are hypothesized to encode information, allowing cells to discriminate between relevant and stochastic signals. However, the mechanisms of frequency decoding and their transcriptional consequences remain incompletely understood. To address this, we investigated how defined Ca²⁺ oscillation frequencies are translated into signaling cascades and gene expression programs in human non-excitable cells. Using optogenetic control of melanopsin-mediated Ca²⁺ influx, we induced slow (8 mHz) or fast (15 mHz) oscillations with identical single-pulse kinetics to isolate the effect of frequency. We found that *TNF* and *IL8* transcription via NF-κB displayed sigmoidal frequency dependence, strictly requiring regular periodic stimulation, while random or low-frequency inputs with equal cumulative Ca²⁺ exposure were ineffective. Bulk RNA sequencing revealed a *MYC*-centered transcriptional response, with 116 of 215 differentially expressed genes predicted as MYC targets, despite unchanged *MYC* mRNA levels. Label-free phosphoproteomics identified PRKDC, CHEK2 and ATM as the top upstream kinases, forming a network linking Ca²⁺ oscillations to cell cycle and stress signaling. These findings demonstrate that cells can decode Ca²⁺ oscillation frequency through a multi-kinase network that tunes transcription via NF-κB and MYC, providing mechanistic insight into how temporal dynamics of second messengers shape cellular decision-making.

## Introduction

Signaling occurs in all cells to regulate intracellular processes and to communicate their state to neighboring cells. Each cell has around 20 distinct intracellular signal transduction pathways with a multitude of interactions [1]. Among these, several are related to Ca^2+^ signaling (for instance cyclic ADP-ribose signaling, voltage/receptor-operated channels and pathways involving activation of phospholipase C), which is involved in a wide array of processes, including fertilization, apoptosis, proliferation, differentiation, muscle contraction, and learning [1-3]. The remarkable versatility of Ca^2+^ as an information mediator stems from its extensive spatial and temporal diversity, allowing it to generate highly varied patterns [4]. Temporarily, Ca^2+^ concentrations can oscillate periodically, driven by the activity of numerous transporters [3]. Modulation of Ca^2+^ oscillatory frequency has been shown to regulate various cellular functions. For example, in the developing brain, ambient GABA and glutamate stimulate interneuron migration. Upon reaching the cortex, upregulation of the potassium-chloride cotransporter KCC2 reduces the frequency of spontaneous Ca^2+^ activity, thereby reducing migration [5]. In post migratory pyramidal neurons on the other hand, the frequency of Ca^2+^ oscillations is increased as the number of branching dendrites increases [6]. Gain-of-function mutations in SHP-2/PTPN11, associated with the developmental disorder Noonan syndrome, enhance the frequency-dependent control of NFAT (Nuclear Factor of Activated T-cells) by Ca^2+^ oscillations, potentially contributing to congenital heart defects [7]. In pancreatic β cells, the frequency of Ca^2+^ oscillations governs mitochondrial ATP synthesis, which in turn drives insulin secretion [8]. Similarly, glioma cells form functional networks and exhibit spontaneous Ca^2+^ oscillations that are fine-tuned to promote proliferation, potentially via NF-κB (Nuclear Factor kappa-light-chain-enhancer of activated B cells) [9]. This activity is similar to the dynamics of developing neural progenitors derived from mouse embryonic stem cells [10, 11].

Biological information encoded in Ca^2+^ oscillations is decoded by various intracellular molecules that adjust their activity in response to these signals [12]. Several molecular mechanisms have been proposed for frequency decoding, with a handful of specific decoders identified to date. These include NF-κB, NFAT, MAPK (Mitogen Activated Protein Kinase), CaMKII (Ca^2+^/calmodulin-dependent protein Kinase II), and calpain [13]. NF-κB and NFAT predominantly decode low-frequency signals in the range of 1-100 mHz, while CaMKII operates in the higher frequency range of 0.1-1 Hz [13]. This separation in the frequency domain is mainly explained by NF-κB and NFAT being shuttled to and from the nucleus as the underlying mechanism of frequency decoding [14], while decoding by CaMKII is dependent on autophosphorylation regulated by Ca^2+^ and calmodulin binding [15].

Optogenetics refers to the use of light to control genetically modified cells that express light-sensitive proteins [16, 17]. Typically, excitable cells such as neurons are activated or inhibited by light stimulation of different versions of the cation channel channelrhodopsin-2 or the chloride pump halorhodopsin. Melanopsin, a G_αq_-protein-coupled receptor encoded by the *OPN4* gene in humans, is expressed in intrinsically photosensitive retinal ganglion cells and plays a critical role in regulating circadian rhythms [18]. Light in the blue spectrum is absorbed by the vitamin A derivate retinal bound to melanopsin, leading to its isomerization from 11-cis-retinal to all-trans-retinal [19]. Upon activation, melanopsin triggers the release of Ca^2+^ from the endoplasmic reticulum (ER) through the inositol 1,4,5-trisphosphate (InsP_3_) receptor (InsP_3_R) [20]. Previously, melanopsin has been expressed in non-excitable cells to enable light-controlled regulation of NFAT activity [21].

In this study, we assay the frequency dependence in the millihertz (period around 1 minute) range of the complete transcriptome and phosphoproteome, in contrast to previous studies that focused on specific decoders to assay their dynamics. We have utilized heterologous expression of melanopsin to precisely control intracellular Ca²⁺ oscillations and to measure downstream frequency decoding in a non-excitable cell line. We analyzed the frequency dynamics of NF-κB and provided evidence for genuine frequency-dependent regulation of the cytokines *TNF* and *IL-8*. We identified a network of frequency-dependent genes centered around MYC through bulk RNA sequencing. Finally, phosphoproteomics revealed a set of proteins with dynamic phosphorylation status, forming a network linking Ca²⁺ oscillations to cell cycle and stress signaling.

## Materials and Methods

### Cell culture

293FT (Invitrogen) and HeLa (ATCC) cells were cultured at 37°C in 5% CO_2_ and regularly passaged. Medium consisted of DMEM (Invitrogen), 10% FBS (Invitrogen) and 1% penicillin-streptomycin (Invitrogen). The day before optogenetic experiments, 100 000 HeLa cells were seeded in six multi-well dishes with or without supplemented all-*trans* retinal (100 nM).

### Light stimulation

A custom-made light stimulation setup was assembled using the following: a blue (470 nm) Rebel LED pre-mounted on a 10 mm square base (Luxeon Star LEDs, MR-B0030-10S) attached to 25 mm square x 10 mm high alpha heat sink (Luxeon Star LEDs, LPD25-10B) using pre-cut thermal adhesive tape (Luxeon Star LEDs, LXT-R-10). Light was focused with a Carclo 18.4 deg lens (Luxeon Star LEDs, 10412). The driving voltage was applied with a power-source (TTi, PL303QMD) connected to a relay (RS, 632-089) and 700 mA constant current LED driver (Recom, RCD-24-0.70/W). The relay was controlled by a DAQ based system (Measurement Computing, USB-1208FS-PLUS) and a custom-made software. Light intensity was measured with a FieldMaxII-TO power meter (Coherent). For studying Ca^2+^ dynamics 5 V was used and for long-term downstream studies 4.3 V was used to reduce light toxicity.

### Calcium imaging

Cells were either loaded with 5 µM Fura-2/AM or 5 µM Fluo-3/AM (both Molecular Probes) in normal culture medium at 37°C for 25 minutes. Then, the cells were washed and immersed in 2 ml of Krebs-Ringer’s medium (in mM): 119.0 NaCl, 2.5 KCl, 2.5 CaCl_2_, 1.3 MgCl_2_, 1.0 NaH_2_PO_4_, 20.0 HEPES (pH 7.4) and 11.0 dextrose. The Ca^2+^ recording experiments were performed using a temperature-controlled chamber (QE-1; Warner Instruments) and a cooled electron-multiplying charged-coupled camera (Quant EM 512 SC, Photometrics) mounted on an upright fixed stage microscope (AxioExaminer.A1, Carl Zeiss) equipped with a 20×/1.0NA objective (Carl Zeiss). Excitation at 480 nm was controlled with an illumination system (DG4, Sutter Instrument). MetaFluor (Molecular Devices) was used to control all devices. Drugs were bath applied.

### Stable cell line

The melanopsin gene (*OPN4*) was cloned into a pLVX-IRES-mCherry expression vector (Clontech PT5061-5) from pIRES_2_-OPN_4_AI for virus production. Lentiviruses were produced by transfecting 293FT cells with pLVX-Melanopsin-IRES-mCherry/pLVX-IRES-mCherry as well as psPAX2 (packaging vector, Addgene: #12260) and pMD2.G (envelope vector, Addgene: #12259) using the Ca^2+^ phosphate method [22]. Thereafter, supernatant was collected and ultra-centrifuged. HeLa cells were transduced with lentiviral particles and polybrene (both with melanopsin gene and only empty vector) and sorted using a BD FACSAria III sorter. Expression was confirmed using RT-qPCR and flow cytometry (BD FACSCanto II).

### RT-qPCR

RNA was harvested using the RNeasy kit (Qiagen) and cDNA was synthesized using the RT^2^ First strand kit (Qiagen). qPCR was performed on an Applied Biosystems qPCR machine using the RT^2^ SYBR Green ROX qPCR Mastermix (Qiagen). Analysis was performed according to the delta delta CT-method [23]. The following primers were used. *OPN4* CCCCTGCTCATCATCATCTTCTG (forward, Tm 67°C) TGACAATCAGTGCGACCTTGGC (reverse, Tm 70°C); *CCND1* TGGAGCCCGTGAAAAAGAGC (forward, Tm 68°C) TCTCCTTCATCTTAGAGGCCAC (reverse, Tm 66°C); *FGF8* GACCCCTTCGCAAAGCTCAT (forward, Tm 68°C) CCGTTGCTCTTGGCGATCA (reverse, Tm 68°C); *BAX* CCCGAGAGGTCTTTTTCCGAG (forward, Tm 68°C) CCAGCCCATGATGGTTCTGAT (reverse, Tm 67°C); *BCL2* GGTGGGGTCATGTGTGTGG (forward, Tm 69°C) CGGTTCAGGTACTCAGTCATCC (reverse, Tm 67°C); *TNF* GAGGCCAAGCCCTGGTATG (forward, Tm 69°C) CGGGCCGATTGATCTCAGC (reverse, Tm 69°C); *MYC* GGCTCCTGGCAAAAGGTCA (forward, Tm 68°C) CTGCGTAGTTGTGCTGATGT (reverse, Tm 65°C) *GAPDH* ACAACTTTGGTATCGTGGAAGG (forward, Tm 65°C) GCCATCACGCCACAGTTTC (reverse, Tm 67°C).

### Luciferase assay

Cells were transiently co-transfected using Effectene (Qiagen) with plasmids carrying NF-κB luciferase reporter and Renilla 24 hours before experiments. Thereafter, cells were harvested, and luciferase activity was measured using the dual-luciferase reporter assay system (Promega).

### Viability assay

Possible toxic effect of light stimulation was assayed using the Annexin V kit (ThermoFisher). Stained cells were analyzed by cytometry (BD FACSCanto II). Positive controls were cells treated with 1 µM staurosporine for 19 hours.

### Bulk RNA sequencing

#### Sample preparation

Cells in six multi-well dishes were washed and enzymatically dissociated in pre-warmed TrypLE Express (Invitrogen). Next, they were counted and diluted to 500 cells/µl and kept on ice all the time. After this, 1 µl of cell suspension was mixed with 1.8 µl of lysis mix with beads (0.09 % Triton X-100, nuclease free water, 11.25 mM DTT, 2.25 mM dNTP and 2 units TaKaRa RNase inhibitor) and incubated in 37°C for 10 min. C1 beads (Invitrogen) were prepared to carry C1-T1-T31. Then, the suspension was mixed and incubated in 72°C for 3 min and cooled to 4°C. Immediately after, 3.6 µl of RT mix (0.98x SuperScript first strand buffer, 6 mM MgCl_2_, 0.81 M Betaine, nuclease free water, 1.6 units TaKaRa RNase inhibitor, 18 units SuperScript II and 2.04 µM C1-P1-RNA-TSO6) was added and incubated in 42°C for 90 min. Then, beads were bond on a magnet and supernatant discarded. 10 µl of PCR mix (111 mM dNTP, 0.48 µM C1-P1-PCR2, nuclease free water and 1x KAPA HiFi hot start ready mix) was added and beads were re-suspended. At last, PCR was run (95°C for 3 min; 6 cycles of 98°C 20 s, 62°C 4 min, 72°C 6 min; 10 cycles of 98°C 20 s, 68°C 30 s, 72°C 6 min; 8 cycles of 98°C 30 s, 68°C 30 s, 72°C 7 min, and 72°C for 10 min) and supernatant saved. The concentration and quality of cDNA was measured using the Quant-iT assay (Invitrogen) and BioAnalyzer (Agilent Technologies), respectively.

#### Library preparation and sequencing

A sequencing library for Illumina HiSeq2000 was prepared using Tn5 transposase. cDNA library was cleaned with AMPure (Beckamn Coulter) and quantified with qPCR. All 32 samples were pooled together and sequenced on one lane.

#### Differential gene expression and principal component analysis

Differential gene expression analysis was conducted using the DESeq2 package in R [24]. A DESeqDataSet object was created from a count matrix and sample metadata with a design formula (∼ Cell type + Stimulation) to account for the effects of cell type and stimulation in the analysis. By stimulation, we mean exposing cells for one hour or 12 hours with one Ca^2+^ transient every minute (∼15 mHz) or every second minute (∼8 mHz). Normalization and differential expression testing were performed, and differentially expressed genes (DEGs) were identified with a false discovery rate (FDR) threshold of < 0.05. To assess overall variability among samples, principal component analysis (PCA) was performed on the RNA-seq data normalized using DESeq2 package. The first two principal components (PC1 and PC2), which captured the largest amount of variance in the data, were selected for visualization and the PCA results were plotted using the ggplot2 package. To assess the relationship between the first few PCs and different variables (Cell type: Melanopsin vs. Empty and Stimulation: no stimulation, 1h and 12h), Kruskal-Wallis tests were used to determine if there were significant differences in PCs between the sub-groups. If significant differences were observed, post-hoc pairwise comparisons were performed using the Wilcoxon rank sum test. P-values from the pairwise tests were adjusted for multiple comparisons using the Benjamini-Hochberg method to control the false discovery rate.

#### Functional enrichment analysis

To elucidate the biological significance of the DEGs, we performed a comprehensive functional enrichment analysis using g:Profiler (version e111_eg58_p18_f463989d, database updated on 25/01/2024) [25]. A list of DEGs with an adjusted p-value threshold of 0.05 was submitted to g:Profiler, to investigate Gene Ontology (GO) and Kyoto Encyclopedia of Genes and Genomes (KEGG) pathway-enriched terms. The analysis was performed using default parameters, except for the option to highlight only driver terms. Terms with a FDR of < 0.05 were considered statistically significant. Upstream transcription factors (TFs) of statistically significant DEGs (adjusted p-value < 0.05) were predicted using the ChIP-X Enrichment Analysis 3 (ChEA3) algorithm (https://maayanlab.cloud/chea3/) [26]. ChEA3 performs TF-target overrepresentation analysis by integrating data from six TF-target set libraries including: GTEx Coexpression, ReMap ChIP-seq, Enrichr Queries, ENCODE ChIP-seq, ARCHS4 Coexpression and Literature ChIP-seq. Predicted TFs were ranked based on their average rank across all TF-target gene libraries (MeanRank). The resulting TF-Gene regulatory networks were constructed and visualized using Cytoscape [27] and made publicly available on the Network Data Exchange (NDEx) platform [28].

#### Gene Set Enrichment Analysis

To investigate the functional relevance of the transcriptomic data, Gene Set Enrichment Analysis (GSEA) was performed using the fgsea package in R (version 1.28.0) [29]. The input for GSEA was the ranked list of all genes from the DESeq2 differential expression analysis, ordered by the Wald test statistic. This approach evaluates gene set enrichment across the entire transcriptome, without applying a predefined threshold for differentially expressed genes [30]. The analysis utilized the Hallmark gene sets (50 gene sets) [31] from the Molecular Signatures Database (MSigDB), which were downloaded as a .gmt file from https://www.gsea-msigdb.org/gsea/msigdb/human/collections.jsp#H (accessed January 10, 2025). The fgsea package was executed with default parameters to calculate enrichment scores (ES), normalized enrichment scores (NES), and associated p-values. The adjusted p-values (FDR) were used to determine statistical significance, with gene sets having an FDR < 0.05 considered significantly enriched. Visualizations, including enrichment plots for significant pathways, were generated using the fgsea and ggplot2 packages in R.

### Phosphoproteomics

#### Preparation of cell lysates

Cell pellets were re-suspended in 150 µl of lysis buffer (8 M Urea, 1% SDS, 50 mM Tris at pH 8.5 and protease and phosphatase inhibitor (Roche)). Cell lysis was done by vortexing and probe sonication on ice to destroy DNA (20% amplitude, 5/2 pulse in 40 s). Lysate was cleared by centrifugation at 4°C, 13k rpm for 10 minutes. Cleared lysate was transferred to new tube.

#### Protein reduction and alkylation

Proteins were reduced with DTT to a final concentration of 5 mM via incubation for 1 hour at 25°C and alkylated with iodoacetamide to a final concentration of 15 mM via incubation for 1 hour at 25°C in the dark. Excess iodoacetamide was quenched by adding 10 mM DTT.

#### Protein precipitation

Methanol, chloroform and water were added with volumes 4:1:3. Washing was done with one volume methanol and a second wash with ice-cold methanol. Pellets were kept between washing steps and centrifugation at 15k rpm for 10 min. Pellets were air-dried (not completely dry).

#### LysC and Trypsin digestion

Protein pellets were dissolved in 8 M Urea and 50 mM Tris at pH 8.5 and diluted with equal volume of 50 mM Tris at pH 8.5 (final urea concentration 4 M). LysC was added (1:50; LysC:Protein) and incubated in a thermo-shaker, overnight with 300 rpm at 24°C. Next, three volumes of 50 mM Tris at pH 8.5 was added (final urea concentration 1 M). Trypsin (1:75; Trypsin:Protein) was added and incubated in a thermo-shaker, 300 rpm at 24°C for 6 hours. Samples were centrifuged at 4°C, 15k rpm for 10 min, pellets were discarded and supernatants saved.

#### Peptide desalting and Phosphopeptide enrichment

Samples were desalted using C18 SepPak (Waters, 50 mg resin, 150 µl column volume, 5% binding capacity) and enriched for phosphopeptides using Pierce TiO2 Phosphopeptide Enrichment and Clean-up Kit (Thermo Scientific^TM^ Pierce^TM^). Manufactureŕs standard protocols were used.

#### PRLC-MS/MS analysis

Phosphopeptides were injected onto the nLC-MS/MS system, Ultimate^TM^ 3000 RSLCnano chromatography system and Q Exactive Plus Orbitrap mass spectrometer (Thermo Scientific). The peptides were separated on a homemade C18 column, 25 cm (Silica Tip 360µm OD, 75µm ID, New Objective, Woburn, MA, USA) with a 120 min gradient at a flow rate of 300 nl/min. The gradient went from 5-26% of buffer B (2% acetonitrile and 0.1% formic acid) in 115 min and up to 95% of buffer B in 5 min. The effluent was electro sprayed into the mass spectrometer directly via the column. A survey MS spectrum was acquired at a resolution of 140 000 in the range of m/z 300-1650. MS/MS data was obtained with a higher-energy collisional dissociation (HCD) at a resolution of 17 500.

#### Peptide identification

Raw files were processed and analyzed with Proteome Discoverer (Ver 3.0, Thermo Scientific). The data was matched against SwissProt Human database (20361 entries, May 2022) using Sequest as a search engine with a precursor tolerance of 5 ppm and a fragment ion tolerance of 0.02 Da. Tryptic peptides were accepted with 0 missed cleavage. Methionine oxidation and Serine, Threonine, Thyrosine phosphorylation was set as variable modifications and cysteine carbamidomethylation was set as fixed modification. Percolator was used for PetideSpectraMatch validation with a strict FDR threshold of 1%. For label-free quantification, the Minora Feature Detector was used with trace length of 5 min, S/N of 1 and max delta RT of isotope pattern multiplets of 0.2 min. A Sequest HT threshold score of 2 was chosen for peptide validation. Only unique peptides were used for relative quantification and proteins were required to pass a protein FDR of 1%.

#### Missing value imputation and principal component analysis

Based on a previous comparative study [32] in label-free proteomics, reporting that random forest–based imputation consistently outperformed other popular methods, missing values were imputed using the missForest R package [33] with 100 trees. To evaluate the overall variability among samples, PCA was performed on the normalized protein and phosphopeptide count data. The first two/three principal components (PC1, PC2 and/or PC3), accounting for the largest proportion of variance, were selected for visualization, and the PCA results were plotted using the ggplot2 package.

#### Differential abundance analysis

Differential abundance analysis of both proteomics and phosphoproteomics datasets was performed using the limma package in R [34]. A design matrix was generated from the imputed, normalized protein and phosphopeptide count data, along with the associated sample metadata, using the model formula (∼ Cell type + Stimulation) to account for the effects of both cell type and stimulation. In this context, “stimulation” refers to one hour of Ca²⁺ transients delivered either once per minute (15 mHz) or once every two minutes (8 mHz). Statistical testing was then conducted, and significantly differentially abundant proteins (DAPs) and differentially abundant phosphopeptides (DAPPs) were identified based on a FDR threshold of < 0.05.

#### Functional enrichment analysis

To explore the biological relevance of the DAPs, functional enrichment analysis was conducted using g:Profiler (version e111_eg58_p18_f463989d; database updated on 25 January 2024). DAPs with an adjusted p-value < 0.05 were submitted to g:Profiler to identify enriched GO terms and KEGG pathways. The analysis was performed with default parameters, except that only driver terms were highlighted. Enrichment results with a FDR < 0.05 were considered statistically significant.

#### Kinase enrichment analysis

Upstream kinases associated with statistically significant differentially abundant phosphopeptides (adjusted p-value < 0.05) were predicted using the Kinase Enrichment Analysis 3 (KEA3) tool (https://maayanlab.cloud/kea3/) [35]. KEA3 identifies kinases whose known or predicted substrates are overrepresented in the input list by integrating multiple kinase–substrate interaction libraries, including experimentally validated and computationally inferred datasets. Predicted kinases were ranked based on their average position (MeanRank) across all available libraries. The HUGO Gene Nomenclature Committee (HGNC)-approved gene symbols corresponding to each phosphopeptide’s protein identifier were retrieved using the biomaRt [36] R package and used as input for KEA3.

#### Retrieval of transcription factor interactors, kinase enrichment analysis and network construction

The top ten enriched TFs at 1h stimulation, predicted by ChEA3 (based on MeanRank; listed in **Table 1**, ChEA3 section), were used as input to retrieve their interactors. “Transcription factor interactors” refer to any molecules reported to physically interact with the given TF in the BioGRID [37] protein– protein interaction database (version 4.4.248, released August 2025; https://thebiogrid.org)). In BioGRID, the search was restricted to *Homo sapiens* and filtered to include only interactions supported by low-throughput (LTP) physical evidence, retaining higher-confidence interactions. Relevant transcription factor interactors are defined here as those interactors shared among at least two of the top transcription factors. Upstream kinases associated with these interactors were then predicted using the KEA3 tool. The resulting Expression-Kinases regulatory network was constructed and visualized using Cytoscape [27] and made publicly available on the NDEx platform [28].

**Table 1.**
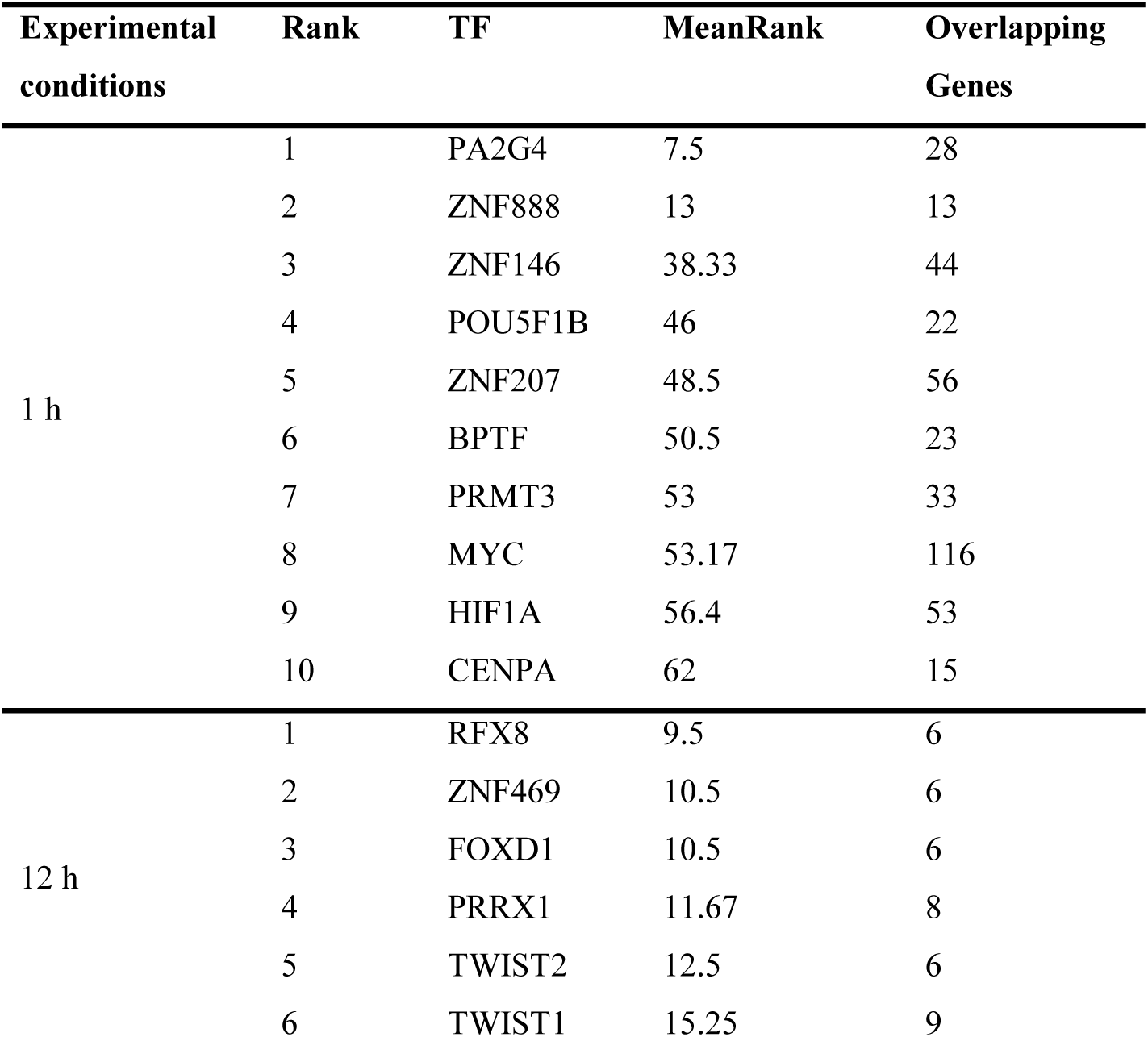
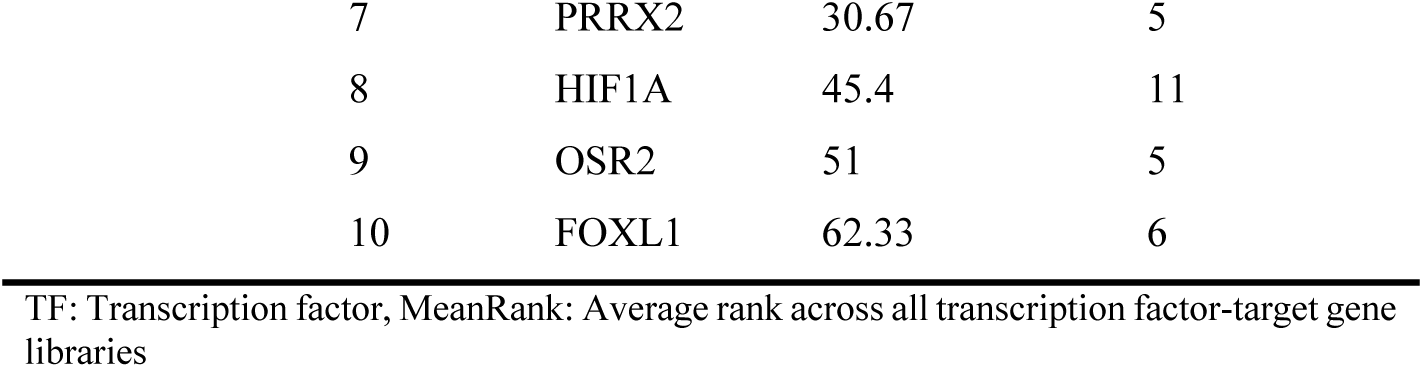
ChEA3 predicted top ten (based on MeanRank) enriched TFs.

| Experimental conditions | Rank | TF | MeanRank | Overlapping Genes |
| --- | --- | --- | --- | --- |
| 1 h | 1 | PA2G4 | 7.5 | 28 |
|  | 2 | ZNF888 | 13 | 13 |
|  | 3 | ZNF146 | 38.33 | 44 |
|  | 4 | POU5F1B | 46 | 22 |
|  | 5 | ZNF207 | 48.5 | 56 |
|  | 6 | BPTF | 50.5 | 23 |
|  | 7 | PRMT3 | 53 | 33 |
|  | 8 | MYC | 53.17 | 116 |
|  | 9 | HIF1A | 56.4 | 53 |
|  | 10 | CENPA | 62 | 15 |
| 12 h | 1 | RFX8 | 9.5 | 6 |
|  | 2 | ZNF469 | 10.5 | 6 |
|  | 3 | FOXD1 | 10.5 | 6 |
|  | 4 | PRRX1 | 11.67 | 8 |
|  | 5 | TWIST2 | 12.5 | 6 |
|  | 6 | TWIST1 | 15.25 | 9 |

|  |  |  |  |
| --- | --- | --- | --- |
| 7 | PRRX2 | 30.67 | 5 |
| 8 | HIF1A | 45.4 | 11 |
| 9 | OSR2 | 51 | 5 |
| 10 | FOXL1 | 62.33 | 6 |

TF: Transcription factor, MeanRank: Average rank across all transcription factor-target gene libraries

## Results

### Optogenetic control of Ca^2+^ oscillations

To study the impact of frequency modulated Ca^2+^ oscillations, we developed a computer-controlled light-setup with optogenetic control of the intracellular Ca^2+^ concentration (**Figure 1A**). The Gα_q_-protein coupled receptor melanopsin, activated by blue light and co-expressed with mCherry, was introduced into HeLa cells. Cells were transduced with lentiviruses with a construct carrying the melanopsin gene *OPN4* co-expressed with mCherry using an IRES sequence (**Figure 1B)**. Simultaneous detection of the intracellular Ca^2+^ level was performed using the ratiometric dye Fura-2/AM. To activate melanopsin, we used a blue LED that emitted light of 470 nm. For live imaging of the kinetics and the influence of the melanopsin receptor on cellular Ca^2+^ levels, we used a setup with one single LED (**Supplementary Figure 1A**), whereas six LEDs connected in series were used (**Supplementary Figure 1B**) for causal studies of frequency modulation of Ca^2+^ oscillations inside the incubator. Various driving voltages were applied to control the radiant power of the LEDs. Measuring the light emitted from one single LED versus one LED connected in series of six revealed similar radiant powers applying 4.3 V and 17 V, respectively (**Supplementary Figure 1C-D**). Testing the response of cells which overexpress melanopsin resulted in a prominent Ca^2+^ increase following 5 s of stimulation with the blue LED (**Figure 1C**). The Ca^2+^ response was dependent on phospholipase C (PLC) and InsP_3_R, since 5 µM of U73122 (**Supplementary Figure 2A**) and 100 µM of 2-APB (**Supplementary Figure 2B**) could inhibit the response. Cells transduced with viruses carrying only the empty vector were not sensitive to the blue LED light (**Figure 1D**).

**Figure 1.**
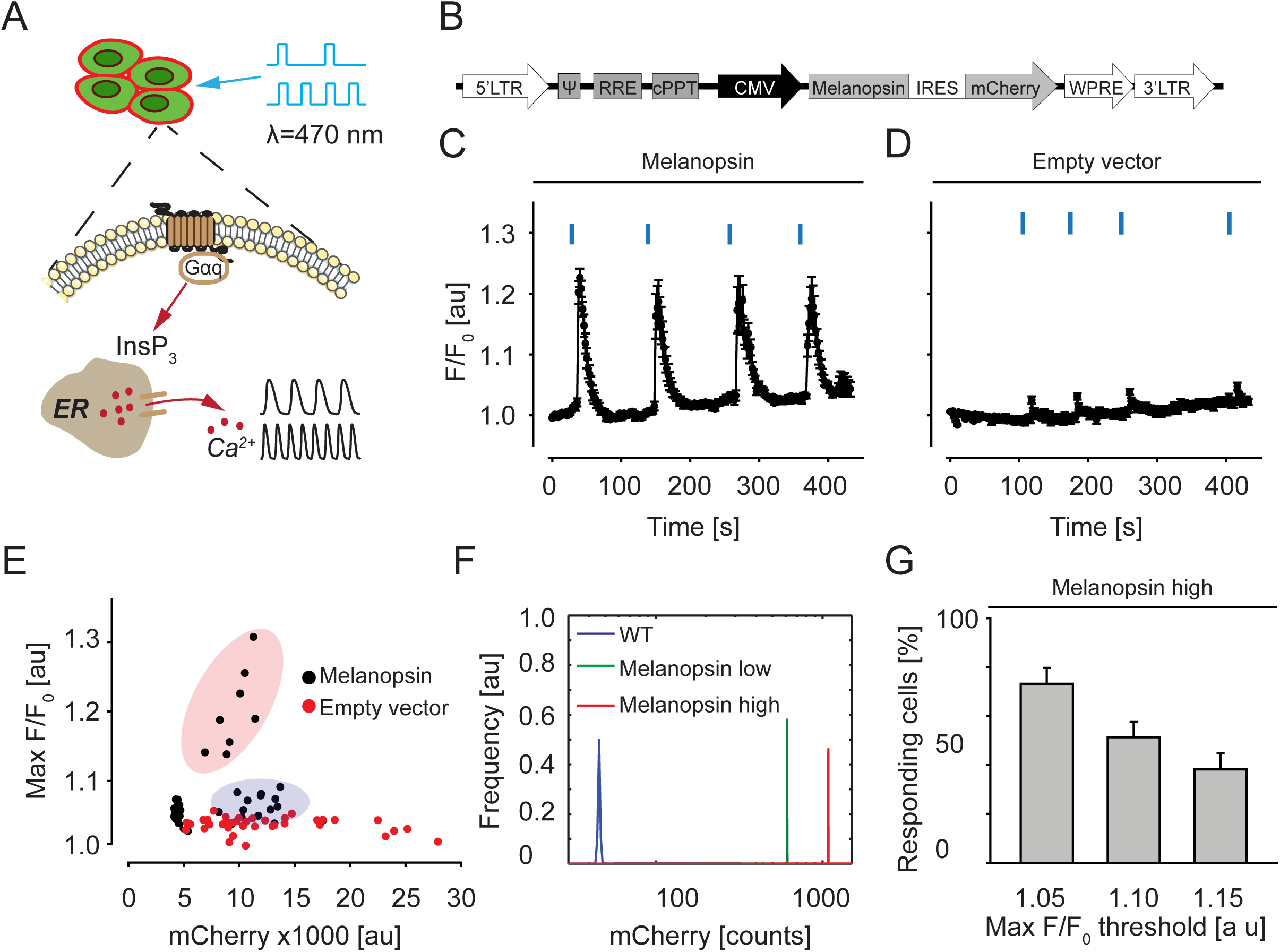
A) Schematic of the experimental setup using optogenetics to control Ca^2+^ signaling. B) The lentiviral expression vector for melanopsin. C) Ca^2+^ trace for the average and SEM for five cells transduced with melanopsin expression vector. Blue markings denote 5 s of light stimulation. D) Same as in C) for cells transduced with empty vector. E) Scatter plot of peak Ca^2+^ response to light stimulation as a function of mCherry intensity. Each dot represents one cell. Red region represents cells with Ca^2+^ response increasing with mCherry fluorescence, whereas blue region represents non-responders. F) Distribution of mCherry intensity for the three sorted cell populations. G) Number of cells responding with Ca^2+^ peaks above threshold, average and SEM for N=3.

Both 5 V and 4.3 V of driving voltage for the LED led to a clear Ca^2+^ response (**Supplementary Figure 2C**). In a population of unsorted cells stimulated with blue light, two different subgroups appeared (**Figure 1E**). In one subgroup (blue region), cells lacked light sensitivity independent on mCherry fluorescence. In the other subgroup (red region), cells with stronger mCherry fluorescence responded with stronger Ca^2+^ influx than cells with weaker mCherry. These data show that we indeed could control the intracellular Ca^2+^ concentration with optogenetics.

### Stable cell lines expressing melanopsin

Next, we sought out to establish stable melanopsin expressing cells. Since there were differences in the Ca^2+^ response depending on the mCherry expression (**Figure 1E**), transduced cells were sorted by flow cytometry into three groups: empty vector, low melanopsin expression, and high melanopsin expression (**Figure 1F**). The mCherry fluorescence corresponded to the melanopsin expression, as validated by RT-qPCR with *OPN4* primers (**Supplementary Figure 2D-E**). To determine if long-term light stimulation was harmful to the cells, we exposed our cells to light-pulses for 5 s every minute for 12 hours. This stimulation was not toxic to cells, as the morphology was unaffected (**Supplementary Figure 2F**) and no increase in the early apoptotic marker Annexin V was observed (**Supplementary Figure 2G-H**). Similar results were observed for cells expressing empty vector, low melanopsin, or high melanopsin.

Thus, neither the light itself nor the resulting Ca^2+^ influx caused apoptosis. In the melanopsin high expressing cell line, around 50% of the cells at a given moment responded with at least 10% increase in Fura-2/AM intensity upon light stimulation (**Figure 1G**).

### Frequency dependence of NF-κB and downstream genes

To validate our experimental system, we first focused on NF-κB, which is known to decode the frequency of Ca^2+^ oscillations. To more specifically study frequency decoding by NF-κB we used a luciferase-based reporter assay. Cells were stimulated with light either once every minute (15 mHz) or once every second minute (8 mHz) for one hour. To test the need for co-factors, culture medium was supplemented in half of the samples with 100 nM of the melanopsin co-factor all-*trans* retinal (ATR). A significantly stronger NF-κB transcriptional activity was found in cells with high expression of melanopsin, as compared to low expression, with ATR (p = 0.0030 and p = 0.020, *N* = 3) or without ATR (p = 0.002455 and p = 0.006597, *N* = 3) (**Figure 2A**). Furthermore, the NF-κB dependent transcription was frequency dependent with higher frequency resulting in stronger expression (**Figure 2B**). The light dependent increase in expression (approximately seven times) was comparable (approximately ten times) to lipopolysaccharide (LPS) dependent activation (**Supplementary Figure 3A**).

**Figure 2.**
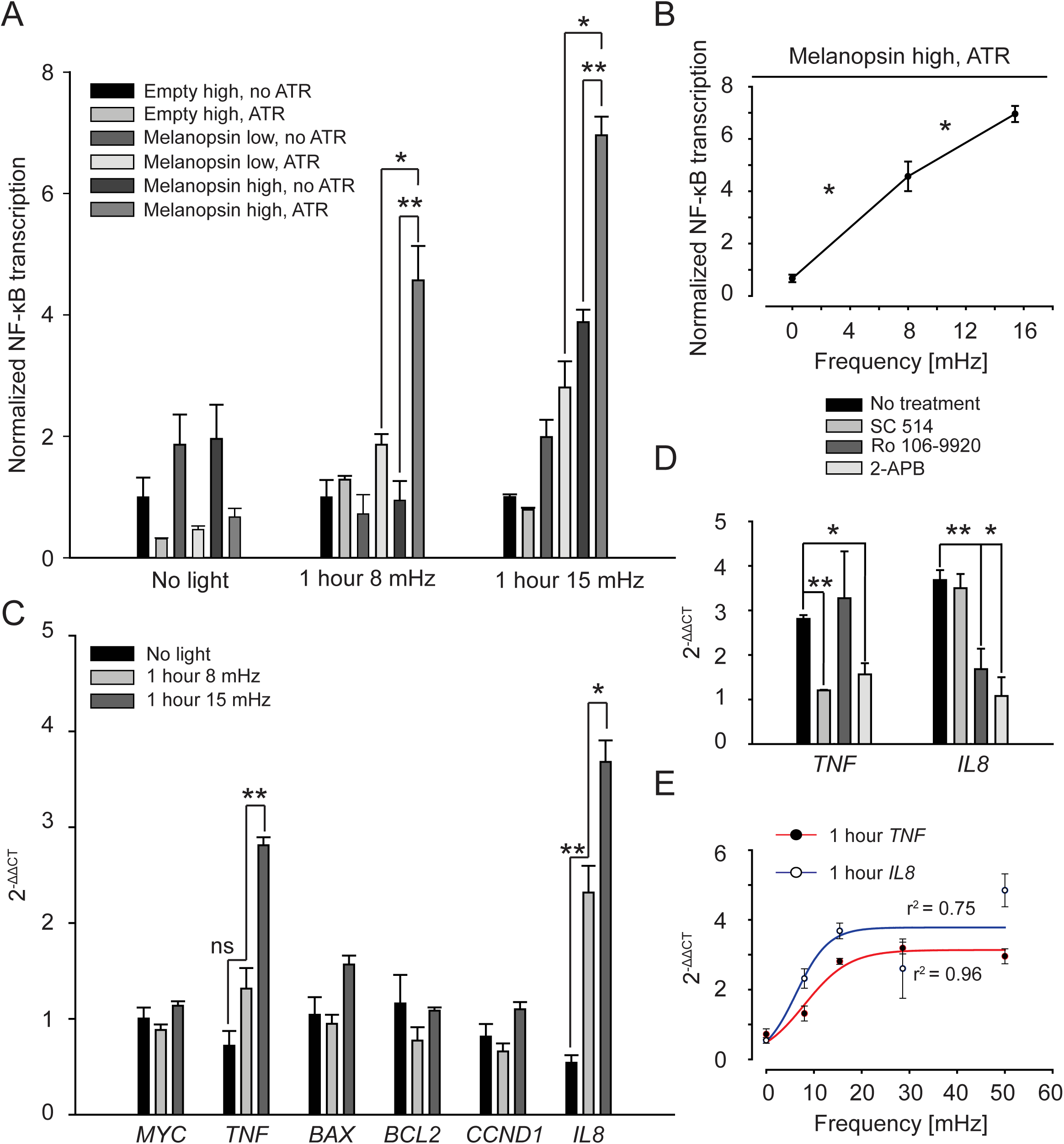
A) NF-κB dependent transcription as measured by luciferase assay. B) Values from A) only for experiments using the Melanopsin high cells with added ATR as a function of frequency. C) Gene expression of a set of genes dependent on NFκB as measured by RT-qPCR. D) The induction of *TNF* and *IL8* is reversed by 100 µM of the IP_3_R blocker 2-APB and the two different NF-κB inhibitors SC 514 (50 µM) and Ro 106-9920 (10 µM). E) Expression of *TNF* and *IL8* as function of frequency for a wider span. P values were calculated with unpaired student’s t test, average and SEM for N=3. * p<0.05 and ** p<0.01.

Next, a set of genes known to be dependent on NF-κB was assayed (*MYC* [38], *TNF* [39], *BAX* [40], *BCL2* [41], *CCND1* [42], and *IL8* [43]) using RT-qPCR. Only the melanopsin high cell line with added ATR was used. The genes *TNF* (p = 0.13868 and p = 0.00607, *N* = 3) and *IL8* (p = 0.00757 and p = 0.03606, *N* = 3) were significantly activated upon one hour of fast and slow Ca^2+^ oscillations (**Figure 2C**). To test if the gene induction was dependent on NF-κB activation, cells were next incubated with either 50 µM of SC 514 (IKKβ inhibitor) or 10 µM of Ro 106-9920 (IκBα ubiquitination inhibitor), which significantly reduced the frequency dependent activation of *TNF* (p = 0.003475, *N* = 3) and *IL8* (p = 0.00211, *N* = 3), respectively (**Figure 2D**). The gene induction was dependent on Ca^2+^ release since addition of 100 µM of the InsP_3_R inhibitor 2-APB blocked the induction of both *TNF* and *IL8* (p = 0.043444 and p = 0.019994, *N* = 3). Surprisingly, cells exhibiting twelve hours of slow or fast Ca^2+^ oscillations did not show significant activation (**Supplementary Figure 3B-C**). To get a more complete picture of the frequency dependence of *IL8* and *TNF,* we then stimulated the cells with a wider range of frequencies. Interestingly, the frequency dependence of the gene expression appeared to be sigmoidal with a clear tendency to saturation for higher frequencies (**Figure 2E**).

### Deeper analysis of frequency dependence of *TNF* and *IL8*

To test if the frequency dependence of *IL8* and *TNF* was dependent on a regular periodicity, cells were stimulated in a random fashion. Cells were exposed to 55 light pulses inducing a random Ca^2+^ oscillatory signal (corresponding to 15 mHz) for one hour (**Figure 3Ai**, lower trace). Only 36% of the pulses were equal to or faster than 15 mHz (**Figure 3B** red region). Intriguingly, the gene induction was significantly weaker with the random stimulation regime compared to the regular (p = 0.022434 and p = 0.010417, *N* = 3, **Figure 3C**). Next, we tested if the frequency dependence of *IL8* and *TNF* was dependent on the exact number of Ca^2+^ increases and/or the elapsed time. Thus, cells were either stimulated every two minutes for one hour (30 pulses, corresponding to 8 mHz), or every minute for half an hour (30 pulses, corresponding to 15 mHz) followed by half an hour without any stimulation (**Figure 3Aii**, upper and lower trace respectively). Interestingly, the gene induction was significantly weaker for *TNF* with the lower stimulation frequency (p = 0.047527, *N* = 3 **Figure 3D**), whereas there was no difference for *IL8* (p = 0.420798, *N* = 3 **Figure 3D**). Together these results conclusively demonstrate that there exists *bona fide* frequency dependence in cells for *TNF* and *IL8*, albeit not completely identical.

**Figure 3.**
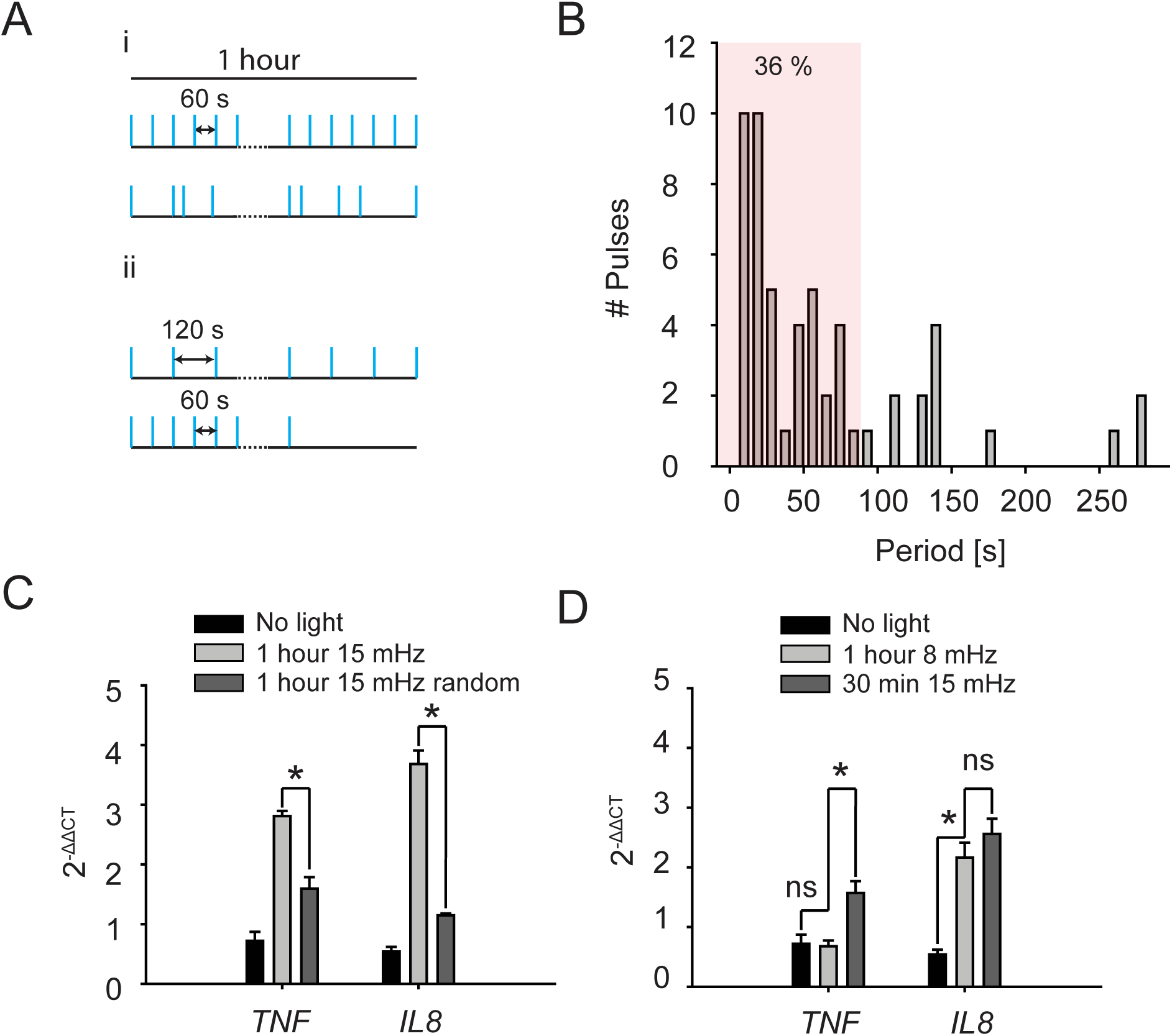
A) Schematic of the two different light stimulation paradigms: i) random stimulation and ii) equal cumulative activity but different frequency. B) The distribution of pause time in-between light pulses for paradigm i). Red region denotes pauses less than 60 s. C) The induction of *TNF* and *IL8* is lower for the random stimulation paradigm i). D) The induction of *TNF*, but not *IL8* is lower for stimulation paradigm ii). P values were calculated with unpaired student’s t test, average and SEM for N=3. * p<0.05 and ** p<0.01.

### Frequency dependence of the full transcriptome

Next, we investigated if light-induced Ca^2+^ oscillations could provoke downstream transcriptional activation in an unbiased fashion. To assay the frequency dependence of the complete transcriptome, we stimulated cells for either one hour or twelve hours with one Ca^2+^ transient every minute (15 mHz) or every second minute (8 mHz). After the experiment, cells were lysed and prepared for RNA-sequencing. In total 32 bulk samples (around 500 cells in each) were multiplexed and sequenced with molecular counting [44].

PCA revealed distinct clustering of samples by cell type (cells expressing melanopsin *vs.* empty vector), with PC1 and PC2 accounting for 84% and 5% of the variance, respectively (**Supplementary Figure 4A**). The separation along PC1 indicates clear transcriptional differences between the two groups. Also considering the type of stimulation (low and high frequency for 1 or 12 hours) reveals clustering as well (**Supplementary Figure 4B**). To assess the relationship between the principal components (PCs) and the Cell_type variable (Melanopsin vs. Empty), Kruskal-Wallis tests were performed. A significant difference was found in PC1 (χ² = 21.05, p = 4.47 × 10⁻⁶), PC4 (χ² = 7.29, p = 0.006933), but not in PC2, PC3, or PC5 (p > 0.05). Assessing the relationship between the PCs and the stimulation variable (Five states), showed a significant association between PC2 and stimulation (χ² = 15.7, p = 0.0034), while PC1, PC3, PC4, and PC5 showed no significant differences across the stimulation conditions (p > 0.05) (data not shown).

The RNA-sequencing analysis identified 215 DEGs between the fast (15 mHz) and slow (8 mHz) oscillating conditions after one hour of stimulation, with “8 mHz” serving as the reference level (**Figure 4A**). Of these, 131 genes were over-expressed, and 84 were under-expressed in the fast-oscillating condition compared to the slow condition. After 12 hours of stimulation, 27 DEGs (17 over-expressed and 10 under-expressed) were identified between the fast and slow oscillating conditions, with the slow (8 mHz) condition again serving as the reference (**Figure 5A**). Of note, there was no overlap between DEGs of 1 hour and 12 hours of stimulation.

**Figure 4.**
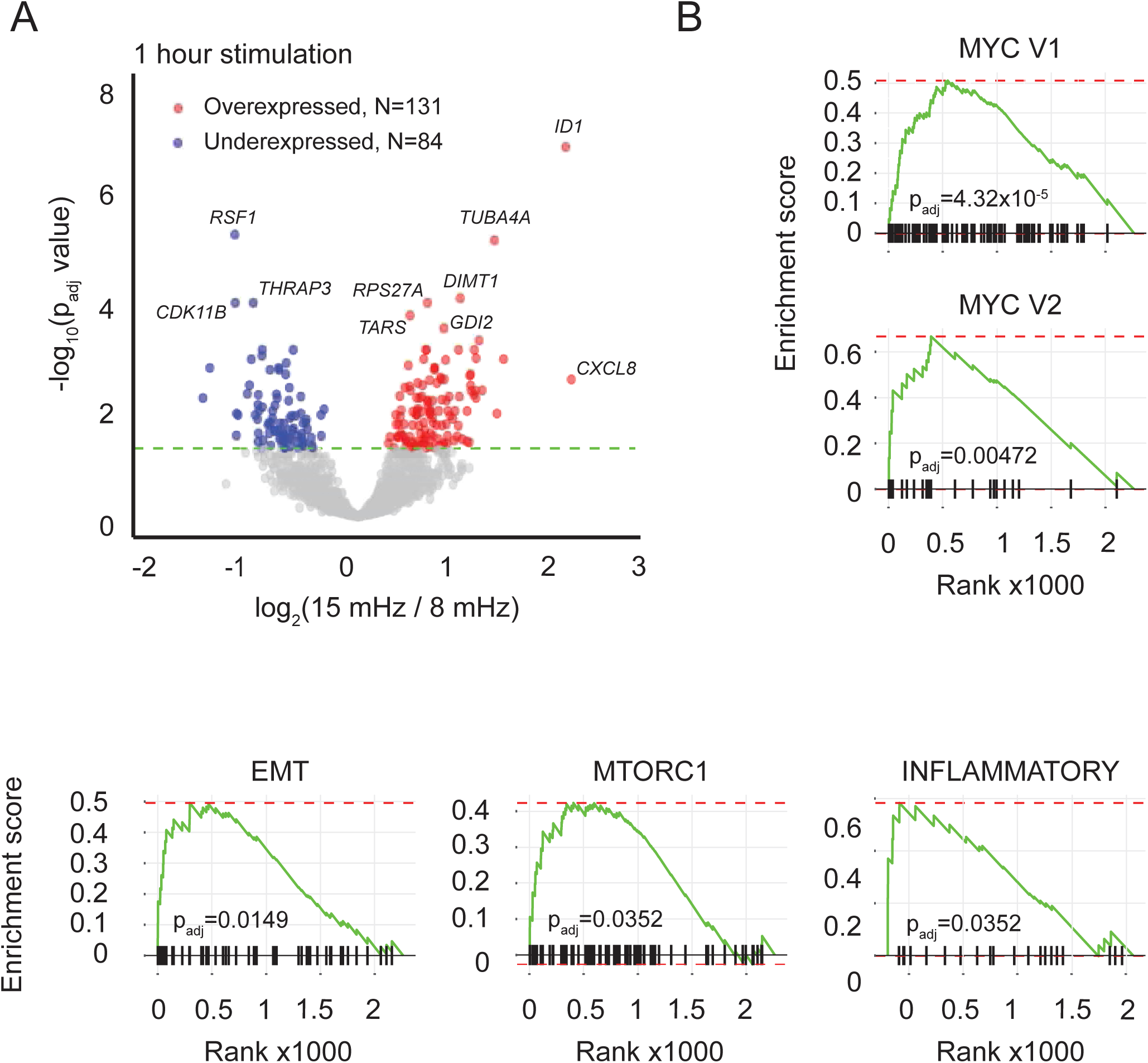
A) Volcano plot of bulk RNAseq data for 1 hour of stimulation showing the log_2_ fold change (x-axis) against the -log_10_ adjusted p-value (y-axis). Genes with adjusted p-value < 0.05 are highlighted: over-expressed genes are marked in red, and under-expressed genes are marked in blue. Non-significant genes are shown in grey. N = 3. B) Enrichment plots of the gene set enrichment analysis results of transcriptomic data. Enrichment plots display the significant pathways identified by GSEA for 1 h using the Hallmark gene sets from MSigDB. The x-axis represents the rank order of genes in the list, while the y-axis indicates the running enrichment score. Tick marks along the x-axis denote the positions of genes from the respective gene set in the ranked list. Pathways with FDR < 0.05 were considered significantly enriched.

**Figure 5.**
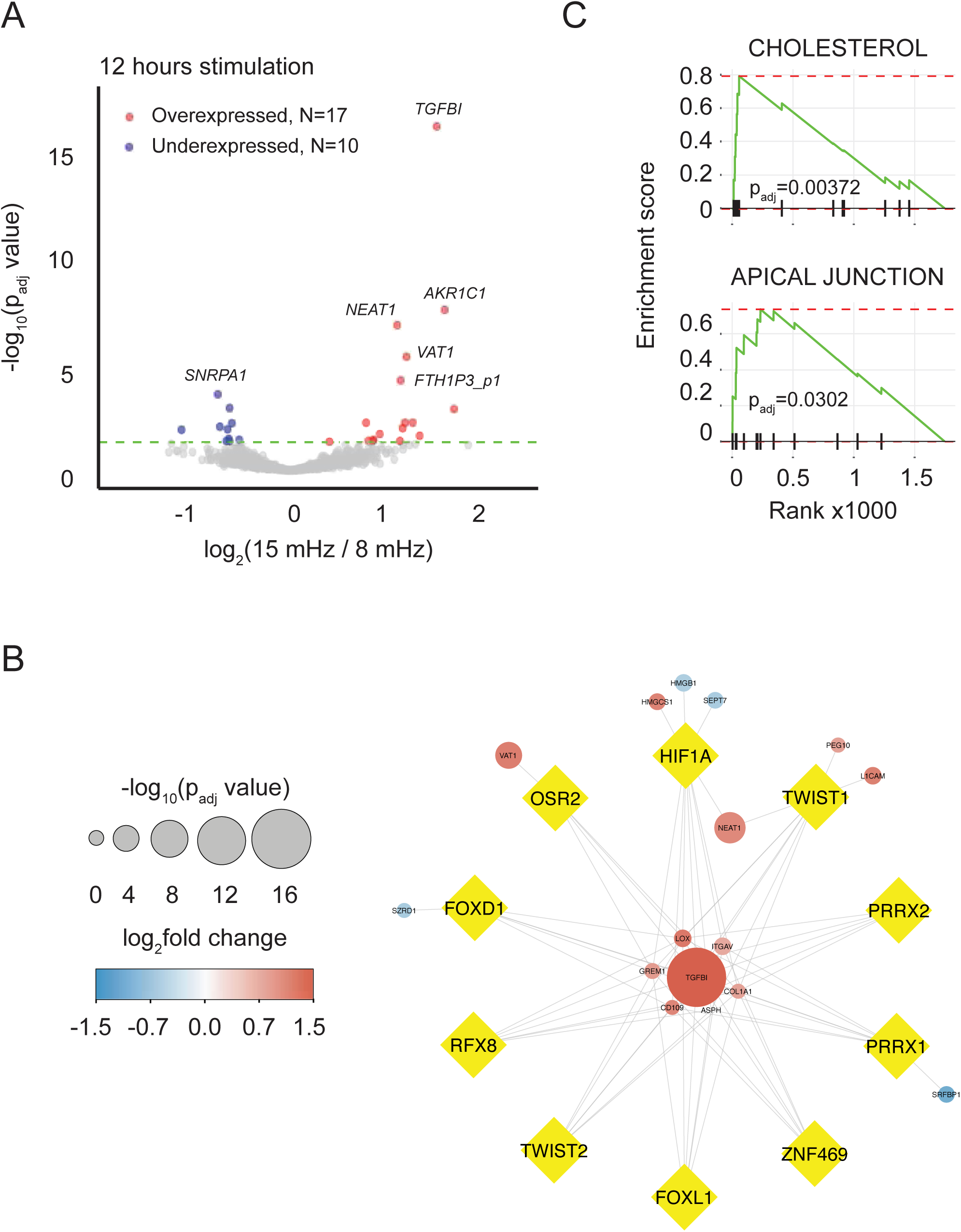
A) Volcano plot of bulk RNAseq data for 12 hours of stimulation showing the log_2_ fold change (x-axis) against the -log_10_ adjusted p-value (y-axis). Genes with adjusted p-value < 0.05 are highlighted: over-expressed genes are marked in red, and under-expressed genes are marked in blue. Non-significant genes are shown in grey. N = 3. B) TF-gene regulatory network constructed using (DEGs) after 12 hours of stimulation. TFs are represented as yellow diamond-shaped nodes, while target genes are shown as circles. Node size corresponds to -log_10_ adjusted p-value, with color representing log_2_ fold change (log_2_FC): red for upregulated genes and blue for downregulated genes. Edges indicate regulatory interactions between TFs and their target genes. C) Enrichment plots of the gene set enrichment analysis results of transcriptomic data. Enrichment plots display the significant pathways identified by GSEA for 1 h using the Hallmark gene sets from MSigDB. The x-axis represents the rank order of genes in the list, while the y-axis indicates the running enrichment score. Tick marks along the x-axis denote the positions of genes from the respective gene set in the ranked list. Pathways with FDR < 0.05 were considered significantly enriched.

GO and KEGG pathway functional enrichment analyses were performed on DEGs using the g:Profiler tool and revealed 26 significant driver terms associated with the 1-hour DEGs. The top three most significant GO molecular function (MF) terms were “protein binding” (GO:0005515), “RNA binding” (GO:0003723), and “protein-containing complex binding” (GO:0044877). The most significant GO biological process (BP) terms included cytoplasmic translation (GO:0002181), ribosome biogenesis (GO:0042254), and organelle organization (GO:0006996). The top three GO cellular component (CC) terms were cytoplasm (GO:0005737), focal adhesion (GO:0005925), and small-subunit processome (GO:0032040).

To identify potential regulators of the Ca^2+^ oscillation-associated gene expression changes, we applied ChEA3, a web-based tool for transcription factor enrichment analysis, to DEGs between fast (15 mHz) and slow (8 mHz) oscillating conditions after 1 and 12 hours of stimulation [26]. The top ten enriched transcription factors for each time point are listed in **Table 1** and shown in **Supplementary Figure 5A and 5B**, respectively.

Subsequently, TF-Gene regulatory networks were constructed for both 1-hour and 12-hour stimulation conditions. The 1-hour network consists of 169 nodes (10 TFs and 159 genes) and 403 edges (**Figure 6**) and is available on NDEx: <u>TF_Gene_Regulatory_Network_1h</u>. MYC regulates the highest number of genes (n=116), followed by ZNF207 (n=56) and HIF1A (n=53). Similarly, the 12-hour network comprises 26 nodes (10 TFs and 16 genes) and 68 edges (**Figure 5B**) and is accessible on NDEx: <u>TF_Gene_Regulatory_Network_12h</u>. For 12 hours, HIF1A regulates the highest number of genes (n=11).

**Figure 6.**
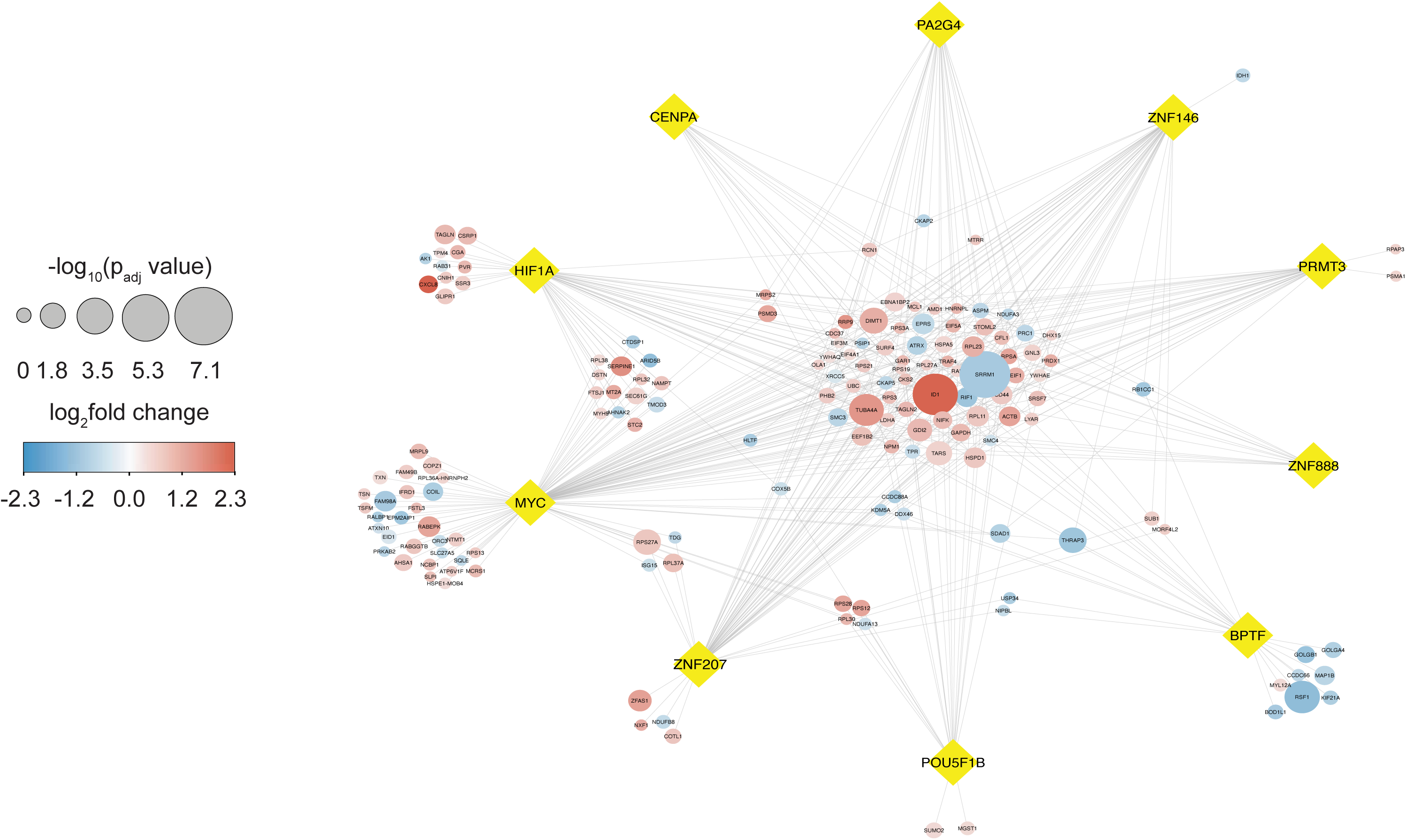
Transcription factor (TF)-gene regulatory network constructed using differentially expressed genes (DEGs) after 1 hour of stimulation under fast (15 mHz) versus slow (8 mHz) Ca²⁺ oscillating conditions. TFs are represented as yellow diamond-shaped nodes, while target genes are shown as circles. Node size corresponds to -log_10_ adjusted p-value, with color representing log_2_ fold change (log_2_FC): red for upregulated genes and blue for downregulated genes. Edges indicate regulatory interactions between TFs and their target genes.

Gene set enrichment analysis was performed to identify significantly enriched pathways associated with the transcriptomic profiles of fast (15 mHz) and slow (8 mHz) oscillating conditions at 1 hour and 12 hours, when taking all genes into account. The ranked gene list derived from DESeq2 was analyzed using the Hallmark gene sets from MSigDB. For 1 hour stimulation, GSEA identified significant enrichment of pathways HALLMARK MYC TARGETS V1 (MYC V1), HALLMARK MYC TARGETS V2 (MYC V2), HALLMARK EPITHELIAL MESENCHYMAL TRANSITION (EMT), HALLMARK INFLAMMATORY RESPONSE (INFLAMMATORY) and HALLMARK MTORC1 SIGNALING (MTORC1) (**Figure 4B**). For 12 hours stimulation, GSEA revealed significant enrichment of pathways HALLMARK_CHOLESTEROL HOMEOSTASIS (CHOLESTEROL) and HALLMARK APICAL JUNCTION (**Figure 5C**). Also, when performing gene set enrichment analysis, there was no overlap in the results between 1 hour and 12 hours of stimulation.

### Frequency dependence of the full phosphoproteome

Next, we assessed the responsiveness of the complete proteome and phosphoproteome to changes in frequency of Ca^2+^ oscillations. Cells were stimulated for 1 hour with light pulses in a similar fashion as for the other experiments and were prepared for phosphopeptide enrichment and mass spectrometry (see **Materials and Methods** for details).

#### Differential abundance analysis of fast *vs.* slow oscillating conditions

Proteomic analysis identified 641 differentially abundant proteins (DAPs) between the fast (15 mHz) and slow (8 mHz) oscillatory conditions after one hour of stimulation, with the 8 mHz condition serving as the reference. Of these, 193 proteins were upregulated and 448 were downregulated in the fast-oscillating condition relative to the slow condition (**Figure 7A**). In the phosphoproteomics dataset, 211 differentially abundant phosphopeptides (DAPPs) were detected, comprising 109 up-and 102 down-phosphorylated peptides, again using the slow (8 mHz) condition as the reference (**Figure 7B**). These peptides corresponded to 92 unique protein identifiers in the up- phosphorylated group and 87 unique protein identifiers in the down- phosphorylated group.

**Figure 7.**
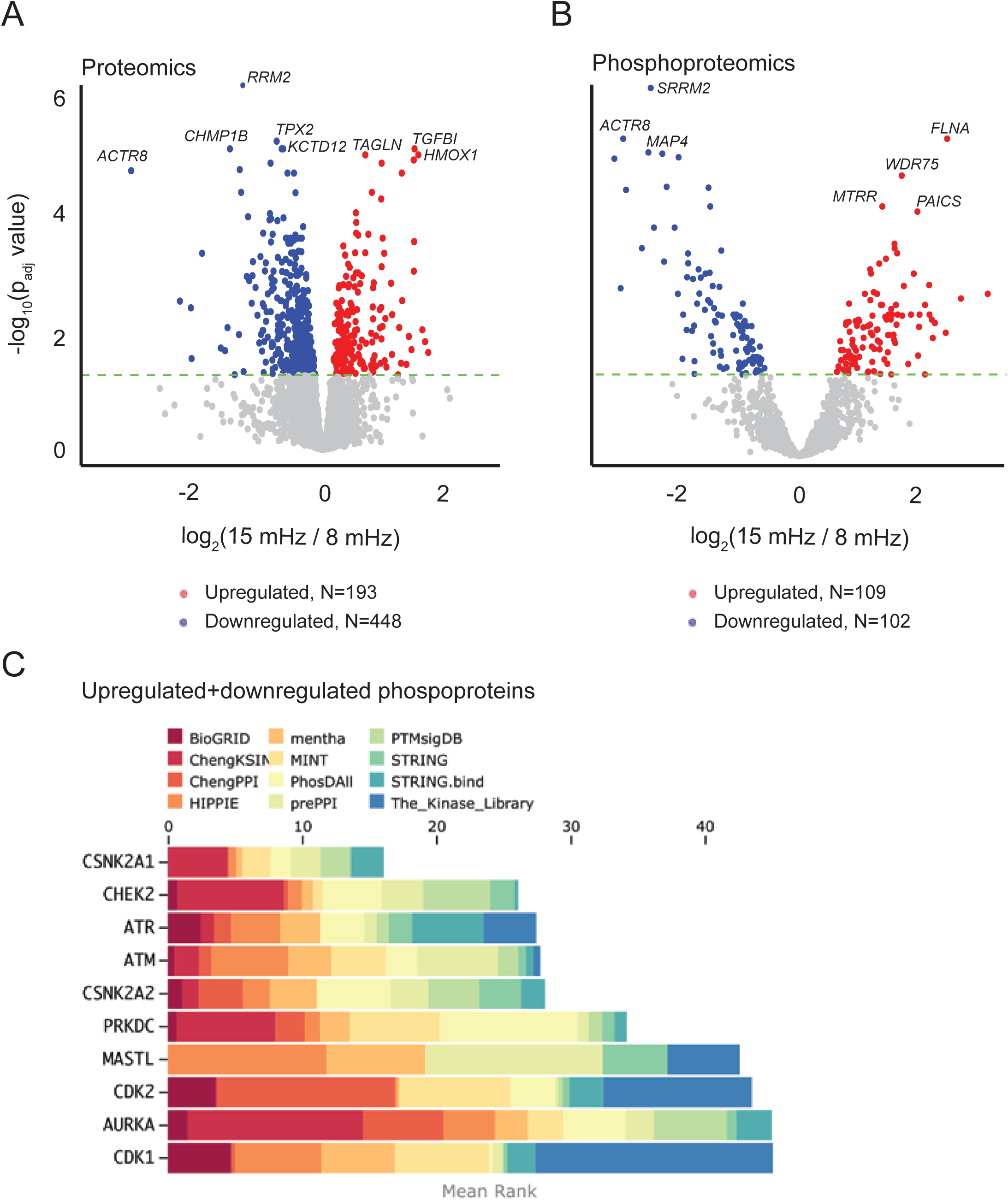
Volcano plots for differential abundance analysis of proteomics and phosphoproteomics at 1 h post-stimulation. Volcano plots for proteomics (A) and phosphoproteomics (B) showing the log2 fold change (x-axis) against the -log10 adjusted p-value (y-axis). Proteins/phosphopeptides with an adjusted p-value < 0.05 are highlighted: upregulated species are shown in red, downregulated species in blue, and non-significant ones in grey. (C) Weighted library contribution to integrated MeanRank kinase ranks. The bar chart showing the cumulative weighted mean rank of the top 10 kinases identified by KEA3 analysis of differentially abundant phosphoproteins. The x-axis represents the cumulative weighted mean kinase rank, while the y-axis lists the top-ranked kinases. The colored segments indicate the contribution of different evidence sources to each kinase ranking.

PCA of the proteomics data using PC1 and PC2 showed limited separation between samples (**Supplementary Figure 4C**); however, inclusion of PC3 improved clustering by cell type (Melanopsin *vs.* Empty vector), capturing additional variance not explained by the first two components (Data not shown). In contrast, PCA of the phosphoproteomics data showed no clear separation between Melanopsin and Empty vector samples when using PC1 and PC2 (**Supplementary Figure 4D**). Inclusion of PC3 in a three-dimensional PCA plot modestly improved the visual distinction between groups, although substantial overlap remained, suggesting that the major sources of variance in the phosphoproteomics dataset are not solely driven by differences in cell type (Data not shown).

#### Functional enrichment analysis of differentially abundant proteins

GO and KEGG pathway functional enrichment analyses of DAPs were performed using the g:Profiler tool. The top three most significant GO MF terms were “protein binding” (GO:0005515), “catalytic activity” (GO:0003824), and “ATP-dependent activity” (GO:0140657). Among GO BP terms, the most significant were mitotic cell cycle process (GO:1903047), organelle organization (GO:0006996), and small molecule metabolic process (GO:0044281). The top three GO CC terms included cytoplasm (GO:0005737), chromosome passenger complex (GO:0032133), and phagocytic vesicle (GO:0045335).

#### Kinase enrichment analysis reveals potential regulators of phosphorylation changes

To identify upstream kinases potentially involved in the downstream Ca²⁺ oscillation–associated phosphoproteomic changes, the consensus sets of up- and down-phosphorylated proteins (unique identifiers) were converted to HGNC gene symbols and submitted to KEA3 tool. The top 10 enriched kinases for up- and down-phosphorylated proteins are listed in **Table 2** and shown in **Figure 7C**.

**Table 2.**
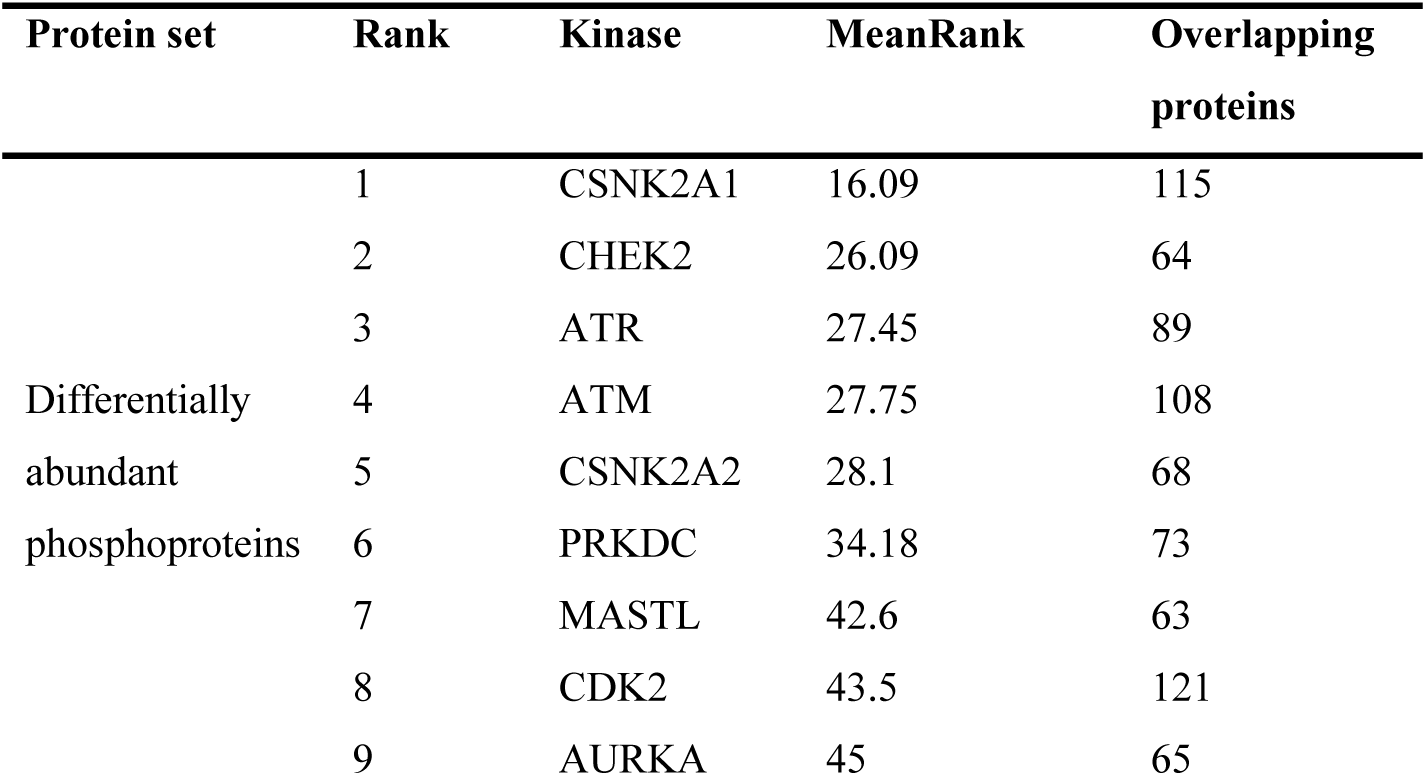
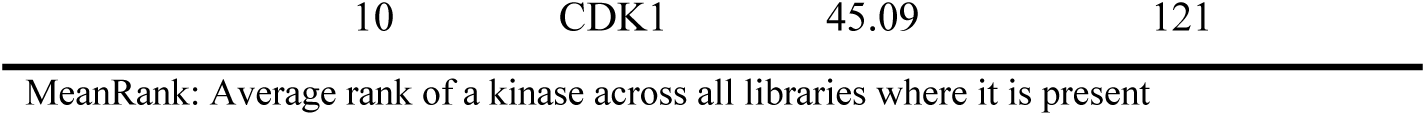
KEA3 predicted top ten (based on MeanRank) enriched kinases.

#### Connecting transcriptomic results with phosphoproteomics

To infer how Ca^2+^ signaling leads to regulation of kinases and downstream transcription factors and gene induction, we sought to combine transcriptomic results with phosphoproteomics. First, proteins potentially interacting with inferred transcription factors were identified. In total, 103 interactors met the minimum criteria and were retained. Of these, 95 were detected on RNA level in the RNA sequencing experiment. This list was used as input to KEA3, and the top ten predicted kinases were retrieved (**Table 3**). Of note, five out of ten predicted kinases were shared between interactors-enriched kinases and those kinases predicted for differentially abundant phosphoproteins: PRKDC (DNA-dependent protein kinase catalytic subunit), CHEK2 (checkpoint kinase 2), ATM/ATR (serine/threonine-protein kinase ATM/ATR) and AURKA (aurora kinase A). Subsequently, Expression-Kinases regulatory network was constructed for 1-hour stimulation condition. The network consists of 109 nodes (8 TFs, 91 interactors and 10 predicted kinases) and 1006 edges (**Figure 8**). The network is also available on NDEx: <u>Expression-Kinases regulatory network</u>. It should be noted that TFs and interactors without direct or indirect connections to kinases were excluded from the network.

**Figure 8.**
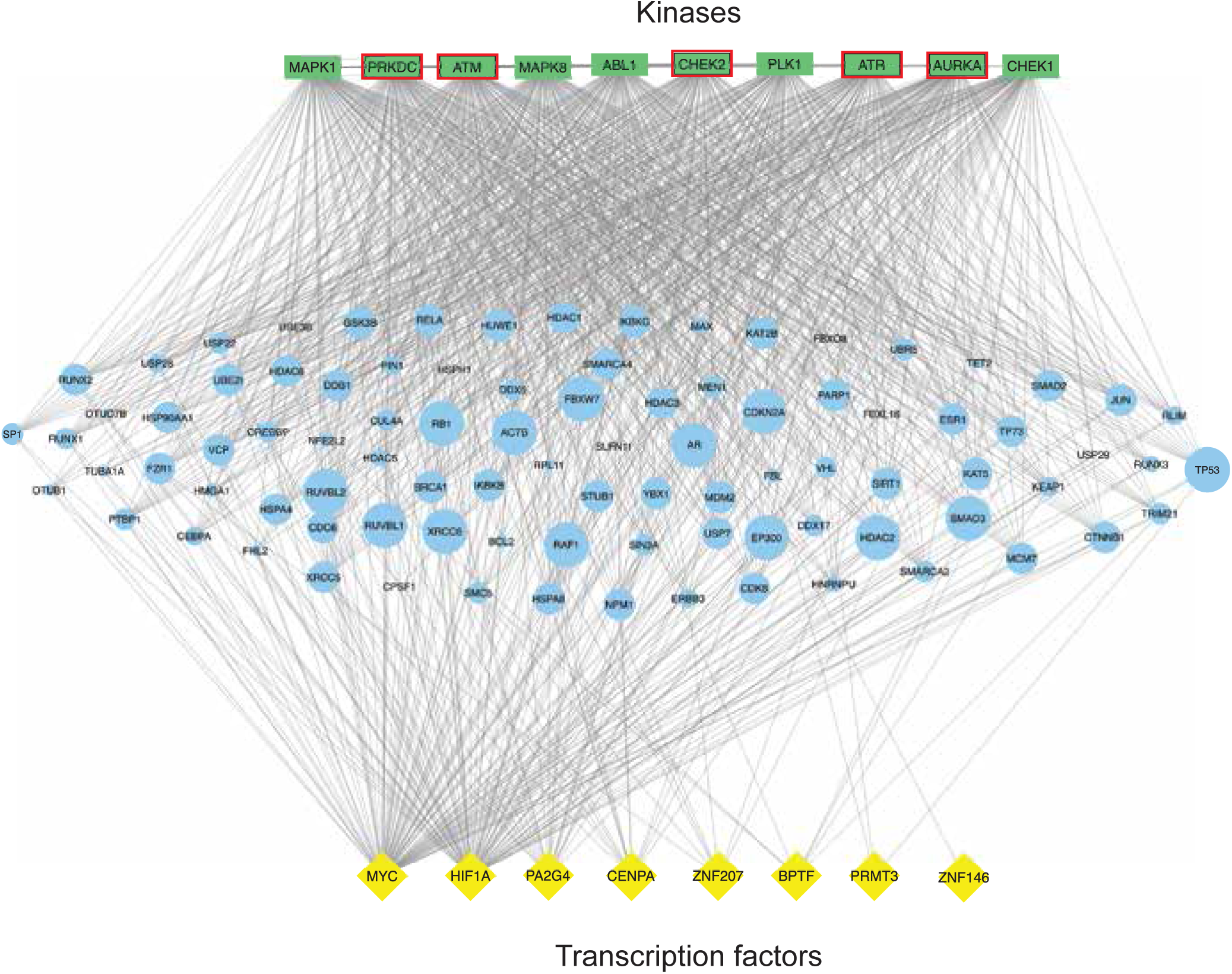
Expression-Kinases regulatory network. The network illustrates interactions between transcription factors (TFs), interactors and kinases. Kinases are shown as green rectangles (top), TFs as yellow diamond-shaped nodes (bottom), and intermediate proteins/transcription factor interactors as blue ovals (middle). The size of interactor nodes is scaled according to their degree (number of connections). Edges indicate regulatory/physical interactions between TFs, interactors and kinases. Kinases with red outlines/borders are those shared between interactors-enriched kinases and ones predicted for differentially abundant phosphoproteins

**Table 3.** KEA3 predicted top ten (based on MeanRank) enriched kinases.

| Protein set | Rank | Kinase | MeanRank | Overlapping proteins |
| --- | --- | --- | --- | --- |
| Interactors | 1 | **PRKDC** | 15.36 | 80 |
|  | 2 | **CHEK2** | 18 | 80 |
|  | 3 | **ATM** | 18.83 | 87 |
|  | 4 | PLK1 | 19 | 80 |
|  | 5 | MAPK8 | 19.09 | 84 |
|  | 6 | CHEK1 | 19.73 | 75 |
|  | 7 | **ATR** | 20.45 | 78 |
|  | 8 | ABL1 | 21.36 | 81 |
|  | 9 | MAPK1 | 26.36 | 88 |
|  | 10 | **AURKA** | 29.27 | 75 |

MeanRank: Average rank of a kinase across all libraries where it is present
Kinases in **Bold** font are those shared between interactors-enriched kinases and kinases predicted for differentially abundant phosphoproteins

## Discussion

Here, we aimed to unravel how intracellular signaling networks decode the frequency of Ca²⁺ oscillations, using optogenetics to precisely control cytosolic Ca²⁺ dynamics. By inducing oscillations with different temporal patterns, we were able to (1) confirm frequency-dependent transcription of *IL8* and *TNF* via NF-κB, and (2) perform an unbiased screening of frequency-dependent signaling using bulk RNA sequencing and phosphoproteomics.

Targeted studies of NF-κB using luciferase reporter and RT-qPCR revealed that both *TNF* and *IL8* exhibit sigmoidal frequency dependence. Importantly, this response was strictly linked to the regular periodicity of the Ca²⁺ signal. Neither random stimulation nor low-frequency stimulation with equal cumulative Ca²⁺ activity elicited an equivalent transcriptional response. This indicates that cells distinguish not just cumulative Ca²⁺ exposure, but also the temporal structure of oscillatory inputs.

RNA sequencing after 1-hour stimulation revealed a MYC-dependent transcriptional signature, supported by both GSEA and ChEA3 analyses. MYC, a bHLH-LZ transcription factor, regulates gene networks controlling proliferation and apoptosis, and its dysregulation underlies many cancers, including cervical cancer (from which HeLa cells originate [45]). While *MYC* mRNA levels were not altered between low- and high-frequency stimulation, its downstream transcriptional network was differentially regulated: of 215 differentially expressed genes, 116 were MYC targets. This suggests that Ca²⁺ oscillation frequency affects MYC activity post-transcriptionally, possibly via phosphorylation or cofactor recruitment. Altogether, 12 hours stimulation led to less differentially expressed genes, potentially due to compensatory mechanisms or less efficient light-induced Ca^2+^ signaling.

Frequency-dependent transcriptional regulation arises downstream of interconnected signal transduction pathways, typically mediated by protein phosphorylation through kinases. To identify the molecular components of this decoding process, we directly measured phosphopeptides using label-free mass spectrometry and inferred candidate kinases involved in frequency-dependent signaling. In parallel, we analyzed differentially expressed genes from RNA sequencing to predict upstream transcription factors and their interacting proteins. Integration of these datasets further allowed prediction of upstream kinases, which showed 50% overlap with those identified through phosphoproteomic analysis when counting the top ten using respective method. Among the kinases inferred from our analyses - PRKDC, CHEK2, ATM, ATR, and AURKA - a unifying feature is their central role in the DNA damage response and cell cycle checkpoint control. These pathways safeguard genome integrity under conditions of replication stress and thereby sustain cellular proliferation. ATM is canonically activated upon DNA double-strand breaks – as well as other threats to DNA integrity – and phosphorylates key proteins, including CHEK2, to inhibit cell cycle progression or induce apoptosis [46, 47]. PRKDC belongs to the same super family of phosphatidylinositol 3-kinase-related kinases as ATM and is involved in the repair of DNA double-strand breaks [48]. Notably, the top three kinases have documented connections to MYC biology. Dysregulation of proliferation and genomic instability in cancers with MYC overexpression is counteracted by ATM and experimental overexpression leads to activation of the ATM DNA damage response pathway [49]. In a similar fashion, inhibition of PRKDC in MYC-overexpressing human lung fibroblasts was shown to be lethal [50]. This convergence suggests that frequency-dependent Ca^2+^ signaling may engage stress-response and checkpoint networks that intersect with MYC-driven transcriptional programs. Interestingly, although ATM, PRKDC, and CHEK2 are not classical Ca^2+^-dependent kinases, there is growing evidence that ATM activity influences Ca^2+^ handling, including mitochondrial Ca^2+^ uptake [51]. Because PRKDC and CHEK2 function downstream of ATM in DNA damage signaling networks, Ca^2+^ dynamics may modulate these kinases indirectly, potentially influencing frequency-dependent signaling responses in our system. Thus, the overlap between kinases inferred from phosphoproteomics and transcription factor analysis may reflect a mechanistic link whereby oscillatory Ca^2+^ signals interface with genome stability pathways to shape MYC-related transcriptional outputs.

Our findings extend the understanding of how temporal coding of Ca²⁺ signals shapes cellular responses. Cells continuously experience a mixture of relevant external cues and stochastic fluctuations, and frequency modulation provides a mechanism to filter noise while preserving information [13]. By directly controlling cytosolic Ca²⁺ with optogenetics, we effectively short-circuited upstream receptor signaling, allowing us to isolate the frequency-decoding capacity of the intracellular network. Previous studies using similar approaches focused on NFAT activation or short-term stimulation [21]. Our study broadens the scope by revealing that frequency modulation affects a wider transcriptional and phosphoproteomic landscape, with MYC emerging as key transcriptional decoder.

Finally, our data addresses the long-standing question of whether cells decode frequency *per se* or cumulative activity. While prior studies on NFAT [21] and NF-κB [52] suggested cumulative activity dominates, we observed that low-frequency stimulation with longer duration did not reproduce the transcriptional response of short, high-frequency stimulation (**Figure 3D**). This indicates genuine frequency decoding under our experimental conditions, although in practice biological systems may integrate both frequency and duty cycle. Using physiologically relevant slow (8 mHz) and fast (15 mHz) oscillations [13] with constant single-pulse kinetics, we ensured that differences reflect temporal coding rather than amplitude variations.

Several limitations of this study should be acknowledged. First, our approach relies on optogenetic activation of melanopsin to directly manipulate intracellular Ca²⁺ dynamics. While experimentally efficient, this strategy bypasses the upstream components of physiological signaling, including ligand binding to surface receptors and the associated cascade of events that recruit Ca²⁺ signaling and kinase activation before transcriptional regulation. Such short-circuiting may alter the fidelity of the responses compared to intact signaling networks. Second, we employed HeLa cells, a non-excitable carcinoma-derived line, as our experimental system. Ca²⁺ signaling toolkits differ across cell types [53], and neuronal and developmental contexts, in particular, possess specialized Ca²⁺ -handling machinery [54]. Thus, the generalizability of our findings to excitable or tissue-specific cell types remains to be established. In addition, the use of label-free mass spectrometry reduces sensitivity for phosphopeptide detection and may therefore underestimate the number of proteins contributing to frequency decoding. Finally, owing to experimental complexity, we did not assess the efficacy of light stimulation in sustaining Ca²⁺ signaling over extended time periods.

In conclusion, our study reveals a comprehensive network of frequency-dependent genes and phosphoproteins using optogenetics, RNA sequencing, and phosphoproteomics. We provide evidence for *bona fide* frequency decoding by NF-κB and identify a MYC-centered transcriptional response potentially modulated by PRKDC, CHEK2 and ATM. These findings open avenues for exploring how frequency decoding shapes cellular decision-making, with implications for signal integration, proliferation, and stress responses.

## Author contributions

ES: designed and performed most experiments, including analysis. Wrote manuscript. PN: performed the RNA sequencing, proteomics and phosphoproteomics analysis. Wrote manuscript. MVG: prepared polyclonal cell lines, including analysis. Wrote manuscript. PU: designed experiments and wrote manuscript.

## Acknowledgments

We are grateful to Dr. Dmitry Usoskin and Dr. Sten Linnarsson for previous development of TTL-based control that was utilized in this project. Dr. Haifeng Ye kindly provided the pIRES_2_-OPN_4_AI plasmid. Dr. Amit Zeisel, Dr. Anna Johnsson and Dr. Marcela Ferella for valuable help with RNA sequencing. Proteomic analysis was carried out at the Proteomics Karolinska (PK/KI) by Carina Palmberg and Dr. Alexey Chernobrovkin as well as at the Proteomics Core Facility, Sahlgrenska academy, Gothenburg University, with financial support from SciLifeLab and BioMS.

---

**SUPPLEMENTARY FIGURE 1.**
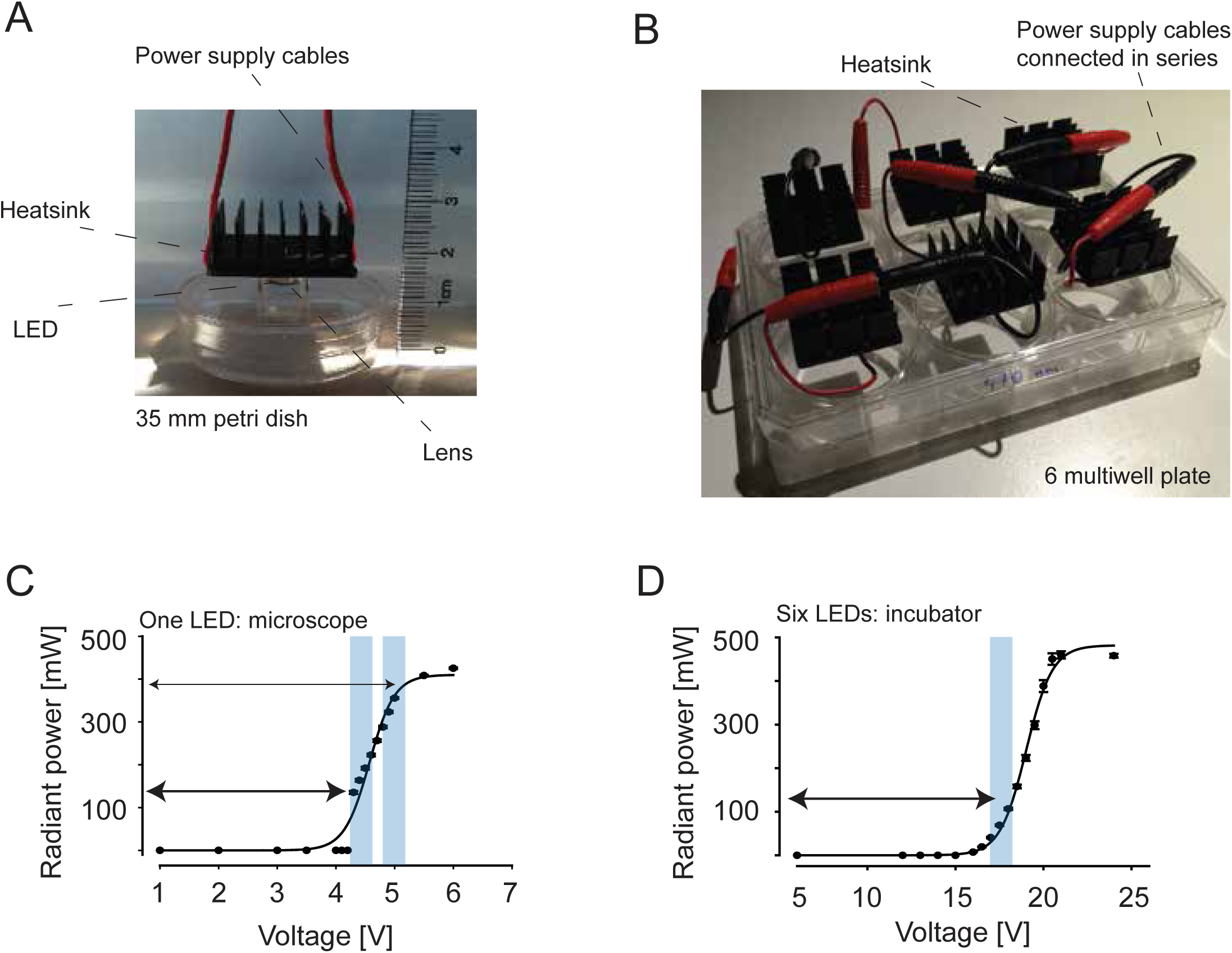

**SUPPLEMENTARY FIGURE 2.**
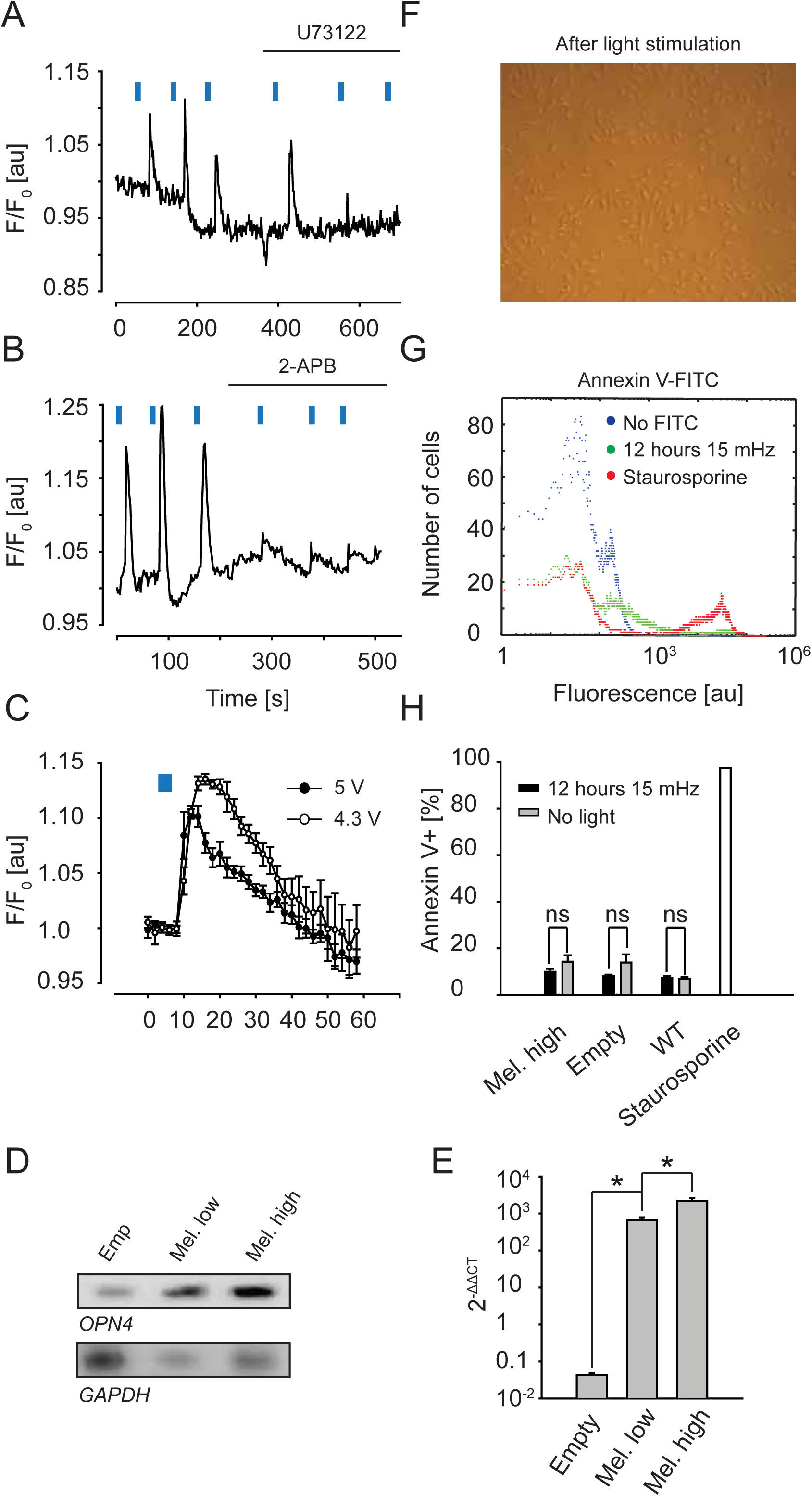

**SUPPLEMENTARY FIGURE 3.**
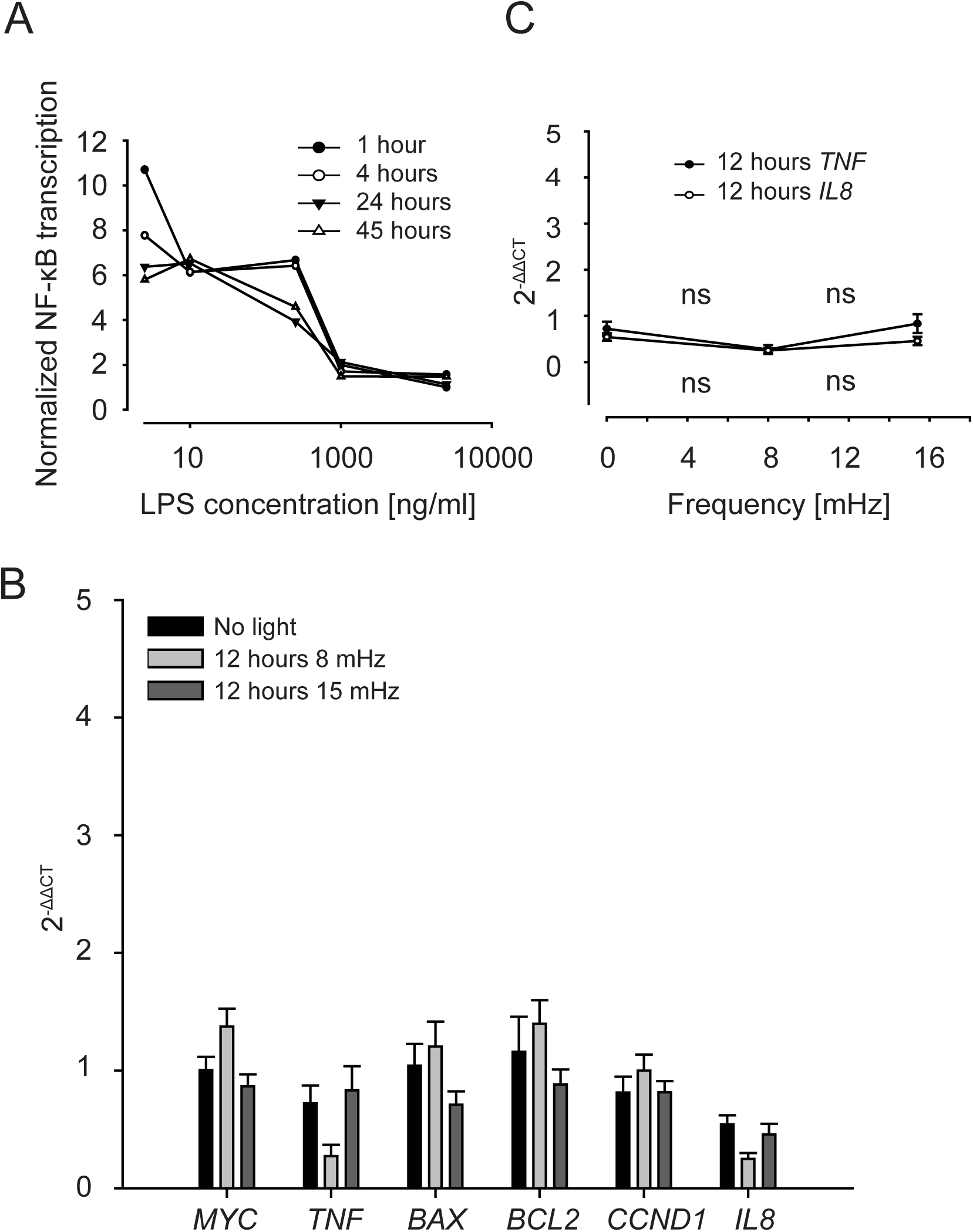

**SUPPLEMENTARY FIGURE 4.**
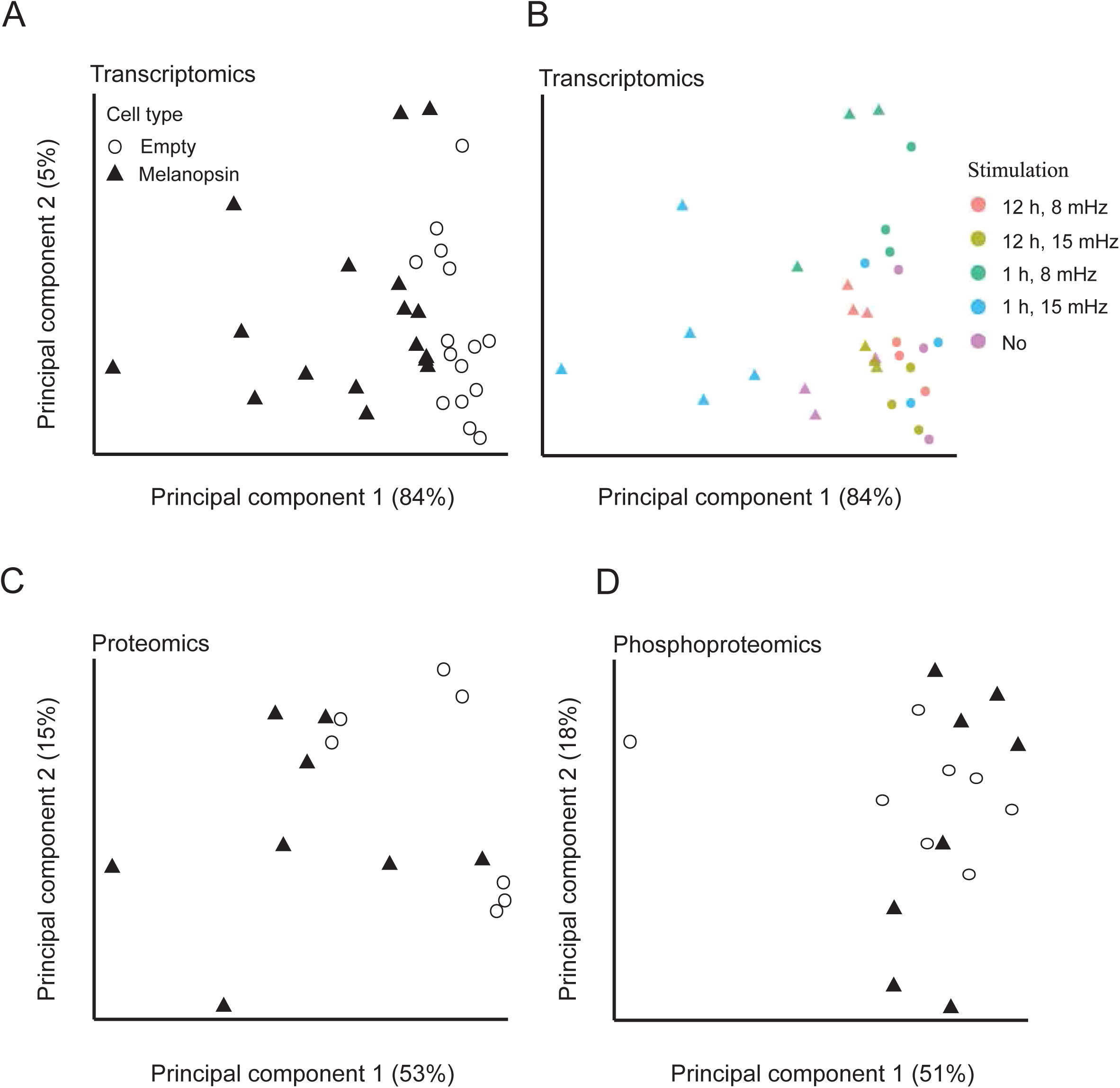

**SUPPLEMENTARY FIGURE 5.**
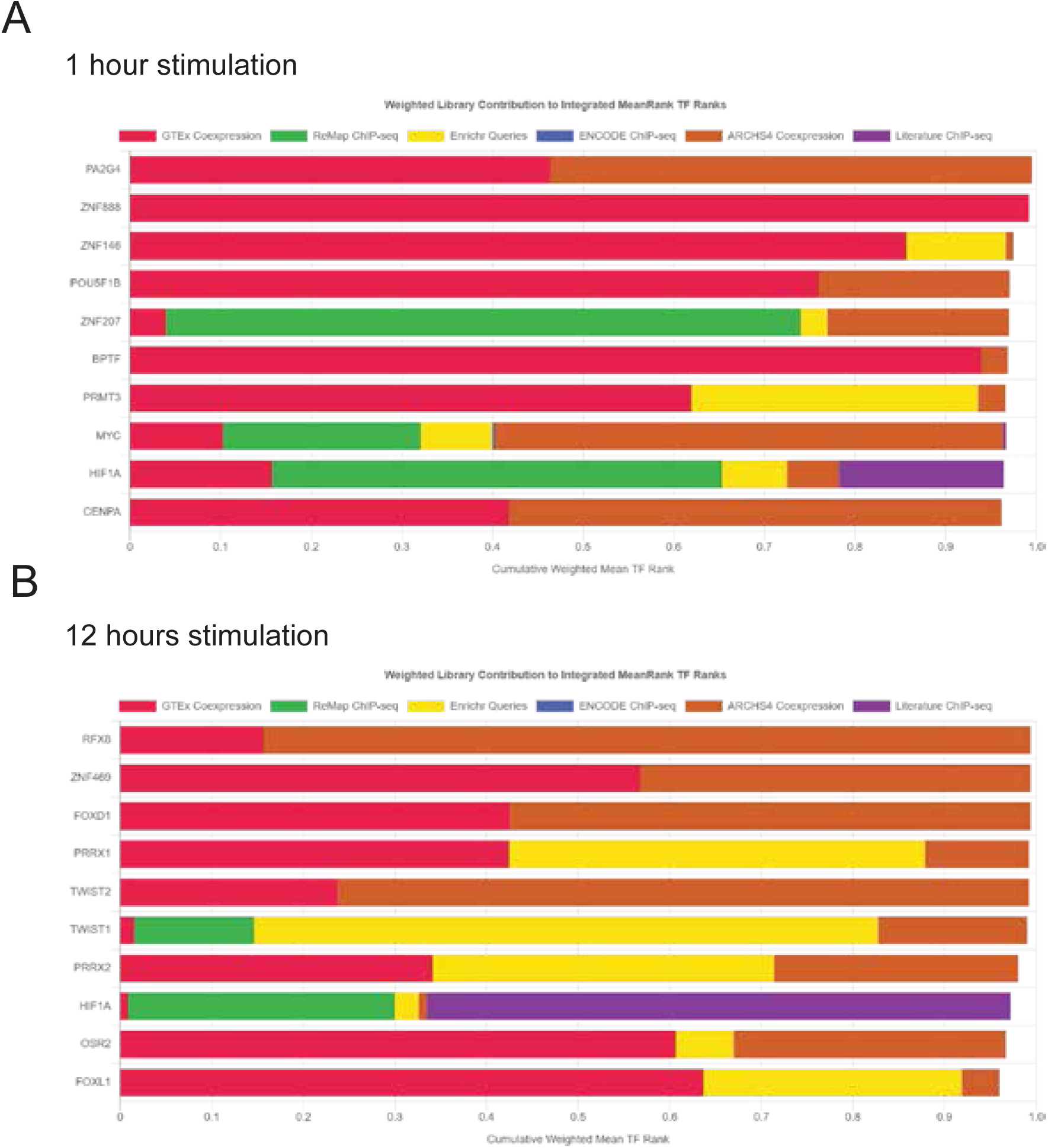

