## Supplementary material for "Decoding calcium oscillation frequency in transcriptional regulation"

### Supplementary figure legends

#### Supplementary Figure 1

A) Picture of LED with cables used for stimulation under microscope and B) in incubator. Light power as a function of driver voltage for C) one LED orD) six LEDs connected in series. Average and SEM for N=3.

#### Supplementary Figure 2

A) $Ca^{2+}$ response to light stimulation is abolished by PLC inhibition with 5 µM of U73122 or B) 100 µM of IP3R inhibition by 2-APB. C) $Ca^{2+}$ response to light stimulation with either low (4.3 V) or high (5 V) driving voltage. D-E) Strong fluorescence corresponds to strong expression of melanopsin as assayed by qPCR. F) Normal cell morphology after twelve hours of light stimulation. G-H) No increase in early apoptotic cells as assayed by Annexin V staining after light stimulation. P values were calculated with unpaired student's t test, average and SEM for N=3. * $p<0.05$.

#### Supplementary Figure 3

A) NF-κB dependent transcription as measured by luciferase assay upon stimulation with LPS at different concentration and duration. B) Gene expression of a set of genes dependent on NFκB as measured by RT-qPCR after 12 hours of light stimulation. C) Expression of *TNF* and *IL8* as function of frequency. P values were calculated with unpaired student's t test, average and SEM for N=3. * $p<0.05$.

#### Supplementary Figure 4

A) Principal Component Analysis (PCA) of RNA-seq samples based on normalized gene expression counts. The PCA plot displays the distribution of 32 RNA-seq samples across the first two principal components, which explain 84% and 5% of the total variance, respectively. Samples are categorized by Cell_type, with circles (○) representing Empty controls and triangles (▲) representing Melanopsin-expressing cells. B) Additionally, samples are grouped by their Stimulation conditions, with the following color scheme: Red for 12 h if 8 mHz, Yellow for 12 h of 15 mHz, Purple for 1 h of 8 mHz, Green for 1 h of 15 mHz and Cyan for No stimulation. (C) PCA of proteomics data projected onto PC1 (52.59% variance) and PC2 (14.91% variance) shows limited separation between groups. (D) PCA of phosphoproteomics data projected onto PC1 (51.19% variance) and PC2 (17.89% variance) shows no clear group separation.

#### Supplementary Figure 5

Bar charts showing the cumulative weighted mean rank of the top 10 transcription factors (TFs) identified by ChEA3 analysis of DEGs after 1 (A) and 12 (B) hours of stimulation under fast (15 mHz) versus slow (8 mHz) $Ca^{2+}$ oscillating conditions. The x-axis represents the cumulative weighted mean

TF rank, while the y-axis lists the top-ranked TFs. The colored segments indicate the contribution of different evidence sources to each TF ranking: GTEx Coexpression (red), ReMap ChIP-seq (green), Enrichr Queries (yellow), ENCODE ChIP-seq (blue), ARCHS4 Coexpression (orange), and Literature ChIP-seq (purple).
